# Chorography and conformational dynamism of the Soluble Human Fibrinogen in solution

**DOI:** 10.1101/2022.09.05.506423

**Authors:** Jose Emiliano Esparza Pinelo, Pragya Manandhar, Grega Popovic, Katherine Ray, Quoc Nguyen, Anthony T. Iavarone, Adam R. Offenbacher, Nathan E. Hudson, Mehmet Sen

## Abstract

Fibrinogen is a soluble, multi-subunit and multi-domain dimeric protein, which, upon its proteolytic cleavage by thrombin, is converted to insoluble fibrin initiating polymerization that substantially contributes to clot growth. The consentaneous structural view of the soluble form of fibrinogen is relatively straight and rigid-like. However, fibrinogen contains numerous, transiently-accessible “cryptic” epitopes for hemostatic and immunologic proteins, suggesting that fibrinogen exhibits conformational flexibility, which may play functional roles in its temporal and spatial interactions. Hitherto, there has been limited information on the solution structure and internal flexibility of soluble fibrinogen. Here, utilizing an integrative, biophysical approach involving temperature-dependent hydrogen-deuterium exchange mass spectrometry, small angle X-ray scattering, and negative stain electron microscopy, we present a holistic, conformationally dynamic model of human fibrinogen in solution. Our data reveal a high degree of internal flexibility, accommodated by four major and distinct flexible sites along the central axis of the protein. We propose that the fibrinogen structure in solution consists of a complex, conformational landscape with multiple local minima, rather than a single, rigid topology. This is further supported by the location of numerous point mutations that are linked to dysfibrinogenemia, and post-translational modifications, residing near fibrinogen flexions. This work provides a molecular basis for the structural “dynamism” of fibrinogen that is expected to influence the broad swath of functionally diverse macromolecular interactions and fine-tune the structural and mechanical properties of blood clots.

## Background

Fibrinogen is a ~340 kDa glycoprotein, which after processing by thrombin, is converted into fibrin and polymerizes into an insoluble scaffold that comprises a major component of blood clots. The soluble form, fibrinogen, consists of a bilateral homodimer with each dimer composed of three polypeptide chains (labelled Aα, Bβ, and γ); the chains are assembled through the formation of five disulfide bridges at the N-terminal ends (Fig. 1A). At 1.5-4 g/L, soluble fibrinogen circulates in the vasculature as one of the most abundant proteins in the blood. [1] Fibrinogen is pivotal in both thrombosis and immunity, and thus, is involved in a range of pathologies from thrombotic to autoimmune disorders such as inflammatory bowel disease, pericellular fibrosis, and atherosclerosis.[1, 2]

**Figure 1.**
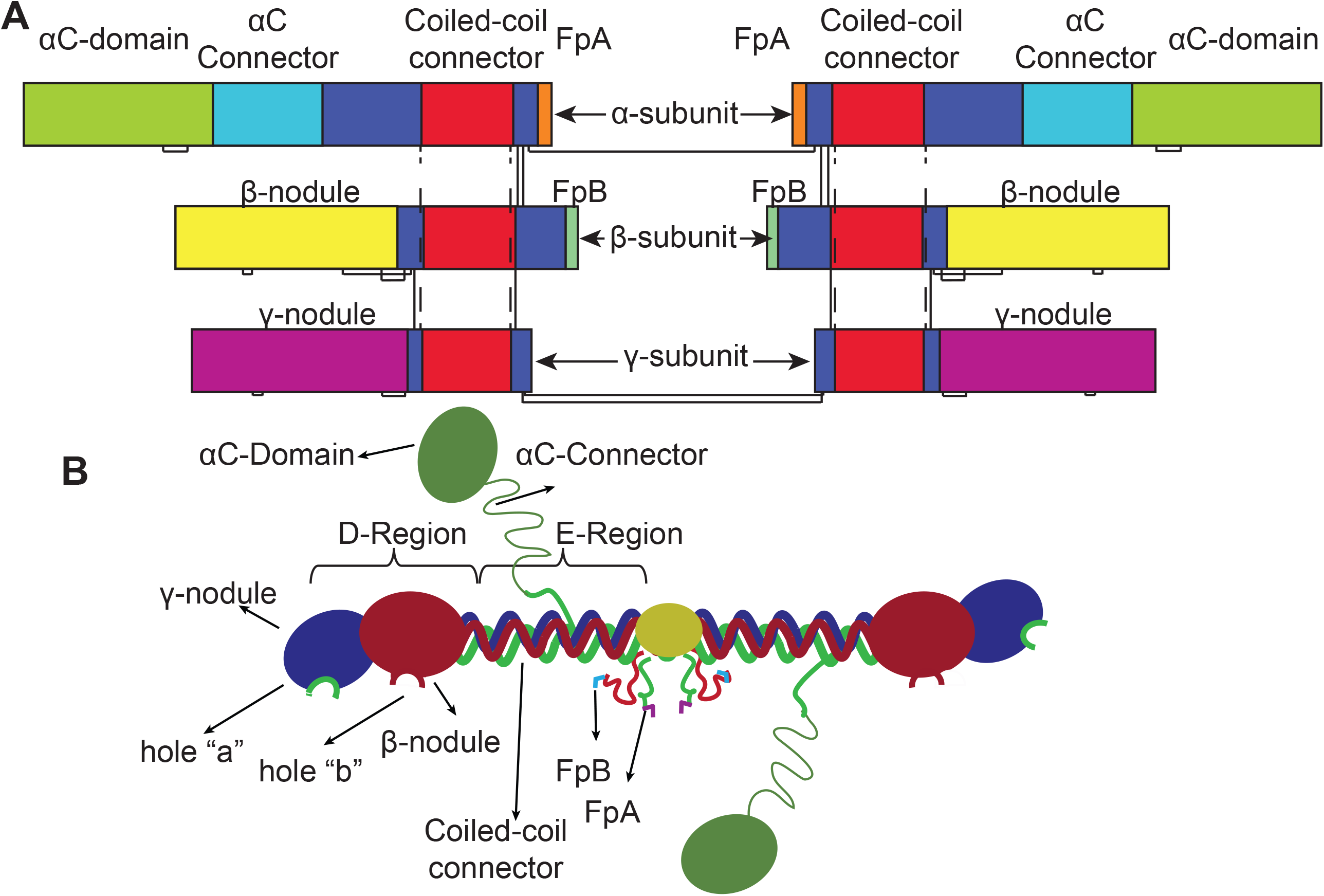
Fibrinogen architecture. **(A)** Domain organization of fibrinogen architecture. Inter- and intrachain disulphide bonds are shown in straight and dotted lines respectively. **(B)** Cartoon representation of the fibrinogen topology. α, β and γ chains are coloured green, red and blue, respectively. Flexion regions detected in this study were shown as red arrows, and functionally important sites and domains are also annotated.

Structural knowledge of the fibrinogen molecule has been amassed over seventy years based on a variety of techniques including electron microscopy (EM), crystallography, and atomic force microscopy (AFM), and the emergent structural model is summarized in Figure 1B. Structural studies showed that the fibrinogen molecule measures ~45 nm end-to-end and resembles a tri-nodular dumbbell. [3–7] From each side of the fibrinogen central nodule, three chains emerge in a 17-20 nm-long coiled-coil helical structure. Disulfide rings flank both ends of the coiled-coil structure providing structural stability and interconnectivity between the chains. The C-termini of the β- and γ-chains consist of globular domains, referred to as the βC- and the γC-nodules. [3, 4, 8] The C-terminus of the α-chain briefly folds back into a fourth coil before extending into a largely unstructured αC connector region followed by a partially-structured αC domain, collectively known as the αC region (Fig. 1B). While the αC region has long been considered to be disordered, investigations into the internal flexibility of central scaffold of the fibrinogen molecule have been experimentally indirect, and thus less compelling.

Early EM-based investigations of the fibrinogen structure reported a mostly straight end-to-end, rod-like conformation, with modest bending in select molecular populations.[9–11] This rigid-like view is consistent with crystallographic images of the full-length human, bovine, and lamprey fibrinogen molecules. A comparison of the crystallographic structures reveals that the dimeric interface about the central nodule is straight, while the coiled-coil region has a slightly twisted, planar-sigmoidal shape where the coiled-coil is bent by a few degrees about its middle (referred to as the coiled-coil connector).[3, 5, 12] Other EM studies, often using solutions without the protein stabilizing agent like glycerol, however, identified heterogenous fibrinogen shapes, which were often bent or curved about the central nodule and/or coiled-coil.[13, 14] Further, the fibrinogen structure studied on a variety of surfaces using AFM has also been observed in bent conformations for a sub-population of molecules.[15–17] More recent, high-resolution AFM-based images also indicate bending of the fibrinogen molecule about its centre and have begun to describe structural aspects of the elusive αC domains.[18] Thus, a myriad of techniques and sample preparations have yielded wildly different views on the conformation(s) and the plasticity of fibrinogen structures. It is important to note that for each of these techniques, the protein is either adsorbed to a surface, or in a crystalline lattice. To interrogate and capture the native conformations of the soluble form of the fibrinogen molecule, it is thus imperative to turn to solution-based techniques.

Due to its high concentration in plasma, fibrinogen is one of the critical determinants of the viscosity and hydrodynamic behaviour of blood. Studies of fibrinogen diffusion and viscosity as a function of concentration, suggest that, in solution, the fibrinogen molecule is much shorter than 45 nm.[19] Combined with molecular dynamics (MD) simulations,[20] these results predicted that fibrinogen molecules explore a much larger “conformational landscape” in solution than hitherto observed in structural studies. A dynamic model of fibrinogen is further implicated by the conformational regulation of fibrinogen’s interactions with various partner proteins; fibrinogen and/or fibrin have binding sites, some of which are cryptic or transiently accessible, for many molecules including plasmin(ogen), tissue plasminogen activator (tPA), integrins [21–23], fibronectin [24], VE-cadherin (CDH5) [25], very low-density lipoprotein receptor (VLDLR) [26], heparin [27], and amyloid-β.[28] However, despite almost identical sequences of fibrinogen and fibrin, there are often differences in the exposure of binding sites for these ligands that distinguish the fibrin(ogen) molecules. For example, plasmin(ogen) and tPA bind a site on fibrin (Aα148-160), that is not exposed on fibrinogen.[29] Similarly, integrin αIIbβ3 binds more tightly to fibrin than fibrinogen [30] but interacts with both fibrinogen and fibrin molecules at distinct binding sites. When interacting with fibrinogen via the γ^406^KQAGDV^411^, αIIbβ3 mediates platelet aggregation, while interacting with the P3-peptide (i.e., γ370-381) of fibrin, αIIbβ3 mediates clot retraction.[31] The RGD sites do not contribute to platelet aggregation, nor do they contribute to clot contraction.[32] Cryptic interactions of the different fibrinogen and fibrin epitopes with leukocyte integrin receptors are also evident.

Aside from modulating the fibrinogen interaction, alteration of the protein’s internal flexibility via single/homozygous mutations or post-translational modifications is expected to play a role in the broad landscape of fibrinogen-mediated clot architecture and phenotypic severity in fibrinogenemia. Several fibrinogen splice variants, as well as an array of post-translational modifications (PTMs), including disulfides, phosphorylation, nitration, and N-/O-linked glycosylation, are linked to (patho)physiological states, such as inflammation and ischemia.[2] In smokers and post-myocardial infarction patients, fibrinogen-nitration—produced by neutrophils and monocytes[33, 34]—carboxylation and oxidation introduce changes in the structural and functional properties of fibrinogen, leading to an increased risk of thrombosis, both arterial and venous.[35–37] Also, hundreds of well-expressed fibrinogen single mutants are associated with three different congenital fibronogenemia.[38] Furthermore, glycan types (e.g., bi- or tri-antennary) and modifications (e.g., de-, mono-, or di-sialylation) on the native fibrinogen have striking effects on the clot architecture. For example, acute phase fibrinogen that is synthesized under inflammatory conditions and fibrinogen that is differently sialylated in neonatal and liver-disease patients showed different clot polymerization kinetics and aberrant fibrin fiber formation.[39–41] Taken together, the cumulative evidence—cryptic binding sites, PTM and single point mutations—implicate that fibrin(ogen) molecules have an intricate shape-shifting mechanism that controls the proper fibrinogen functional output.

Biophysical studies that elucidate the postulated [20] *in solution* dynamics and conformational ensembles would provide a more physiologically-relevant, topological description of fibrinogen and therefore uncover deeper insight into the molecular mechanisms that tune and regulate fibrinogen-dependent events in health and pathologies. Herein, we present a holistic structural and dynamical model for the in-solution, soluble form of human fibrinogen developed through an integrative approach using a combination of temperature-dependent hydrogen-deuterium exchange mass spectrometry (TD-HDXMS), small angle X-ray scattering (SAXS), and negative stain electron microscopy (EM). Our results from SAXS and negative stain EM support a heterogenic dynamic ensemble of fibrinogen conformations, many of which are ‘obtuse-bent’ and asymmetric along the coiled-coiled axis and synchronously moving the βC- and γC-nodules relative to the coiled-coiled axis. TD-HDXMS reveals a “breathing” patch located along the coiled-coil and proximal to the predicted αIIbβ3-integrin interaction site, α-^95^RGD^97^, and an *N*-linked glycosylation sequon, γ-^52^NXS^54^[40] of fibrinogen. The cumulative data herein present a new dynamic view of fibrinogen and functional implications of the fibrinogen structural dynamism in the context of polymerization, molecular recognition and mutation-linked pathologies.

## Results

### TD-HDX Spatially Reveals a “Hinge” in Coiled-Coil Region

First, we used TD-HDXMS to describe the regional protein flexibility of the complete soluble form of fibrinogen. From our pepsin digests, tandem LC-MS/MS supported the assignment of 170, 101, and 119 peptides from the α, β, and γ chains of fibrinogen (SFig. 1-3), providing coverage of 70, 81, and 74% the primary sequence, respectively. Most notable was the large gap in coverage of the α chain, residues 250-347, that represents the αC-connector, a flexible linker between the αC and coiled-coil domains (SFig. 1,4). This αC-connector was likewise absent in previous HDX-MS analysis.[42] This may be attributed to the highly dynamic properties expected for this linker region. In addition, a significant portion of the uncovered regions in α (9-53), β (42-86), and γ (10-28) chains represent the compact region of the central domain which is connected through several post-translational disulfide linkages. This compacted, cysteine-rich region, however, is well structured in the X-ray crystal structure and is anticipated to have minimal amide hydrogen exchange with D_2_O. Further, in the N-terminus of the α chain, there are phosphoserine PTMs that might prevent peptide assignments in this region. Two additional regions, 361-374 in β and 44-56 in γ, were also uncovered, both of which contain N-linked glycosylation sequons, β-N364 and γ-N52.

From the 390 total peptides assigned from the pepsin digests of human fibrinogen, a set of 23 (α), 23 (β) and 21 (γ) non-overlapping peptides were selected for data reduction purposes. Continuous labelling hydrogen exchange was carried out at 10 and 25°C with buffered deuterium under which EX2 conditions are expected and confirmed based on the shift of the mass centroid of the isotopic envelope. Temperature dependent HDX is becoming more prevalent for its ability to provide additional insight into the role of protein fluctuations and their relevance to function, over conventional HDX analysis at a single temperature. [43–45]

Based on pattern recognition, three classes of temperature dependent HDX behaviour emerged (Fig. 2A-C). Class I (Fig 2A) is characterized by low exchanging behavior (<40%) that exhibits no significant differences in HDX between 10 and 25°C. Class II (Fig 2B) is characterized by moderate exchange accompanied by temperature-sensitive apparent rates of exchange and/or extent of exchange (i.e., protein motions are thermally activated). Finally, class III (Fig 2C) is characterized by high exchange behavior (>60%) that likewise exhibits no significant differences in HDX between the two temperatures. The latter behavior is expected for peptides that are either unstructured and/or highly dynamic in nature.

**Figure 2.**
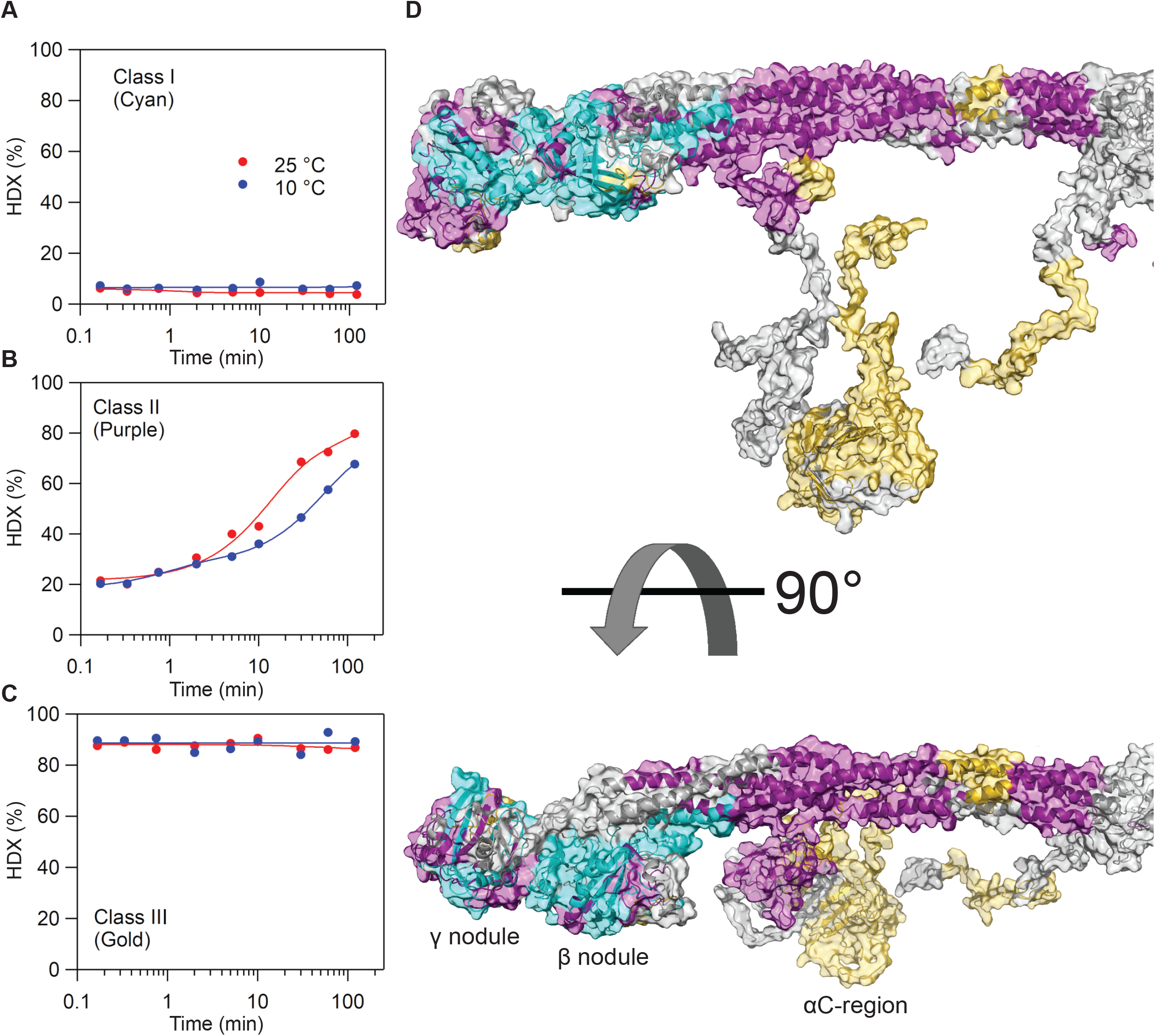
Classification of TDHDX-MS properties of human fibrinogen. Representative traces of exchange properties of fibrinogen-derived peptides were assessed at two temperatures, 10 (blue) and 25°C (red) (A-C). (A) Class I corresponds to low, temperature-independent exchange. (B) Class II corresponds to moderate, temperature-dependent exchange. (C) Class III corresponds to moderate-to-elevated, temperature-independent exchange. In (D, E), the color coding, corresponding to the three classes, is mapped onto the fibrinogen model. Uncovered regions are represented by gray coloring.

We mapped the exchange classes onto the fibrinogen structural model (Fig. 2D, E). From this view, class I is almost completely centered on the β- and γ-nodules, supporting that these regions of the protein are rigid and well structured. Class III is mostly restricted to the αC region of fibrinogen, with high extents of exchange (>80 %). These data support the hypothesis that the αC-domain is highly flexible and potentially shows “breathing”-like characteristics. Conversely, the coiled-coil is almost predominantly characterized by class II (Fig 2D, purple) behavior. One exception to this behavior is a short sequence (α:74-89 and β:106-115) that exhibits temperature-independent, high exchange consistent with class III behavior. Our TD-HDX data support that this localized “hinge” region is highly dynamic and could allow flexing of the fibrinogen molecule as previously predicted from MD simulations.[20] To further visualize the hinging motions of fibrinogen, we also pursued structural characterization of fibrinogen by both EM and SAXS.

### EM of Fibrinogen Reveals Structural Heterogeneity

Intact fibrinogen was examined by negative stain EM in 150 mM NaCl and 20 mM HEPES pH 7.4 at 1.26 Å/pixel and 5.46 Å/pixel resolutions. In high-resolution images, although most fibrinogen particles were elongated and asymmetrical, the general dimeric features of the βC- and γC-nodules were easily discerned (Fig. 3A). In some single particles, the globular αC-domain protrusions were evident (Fig. 3A-red arrows). To further categorize the overall conformations, more than 3000 particles were picked and then, images were subjected to 10 cycles of multireference alignment and *K*-means classification into 50, 20 and 10 sub-classes using SPIDER (SFig. 5).[46] Image averages revealed heterogeneous conformational ensembles, described by predominantly obtuse bends along the major axis (SFig. 5B and C).

**Figure 3.**
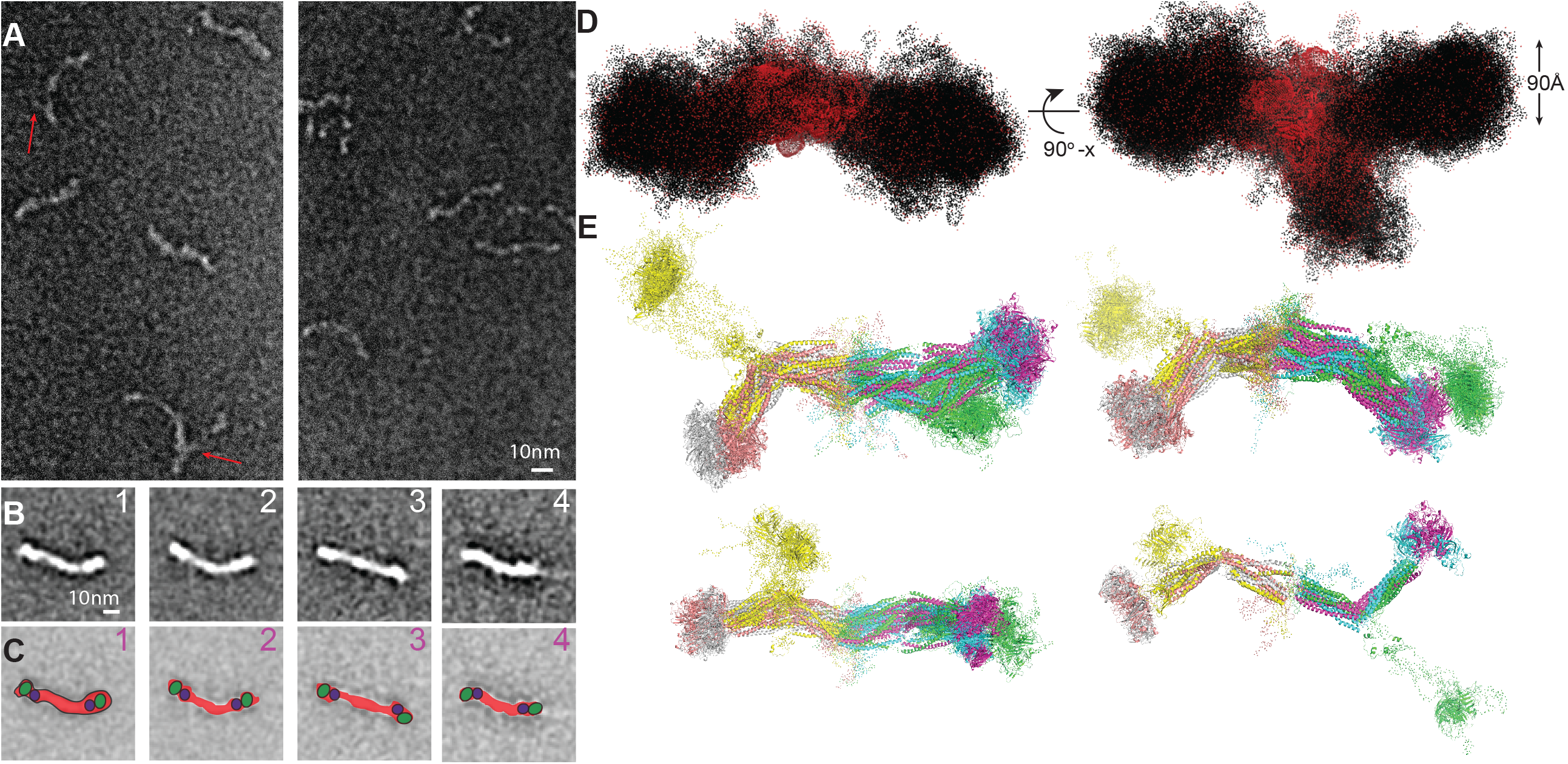
Negative Stain EM and SAXS structures of fibrinogen. **(A)** Raw images of negatively stained fibrinogen. The αC-domain protrusions observed in some of the particles are pointed with the red arrows. **(B)** The four most frequent class averages and **(C)** their 2D projection shadows that correlate best to class averages in panel B. The red, green and purple areas represent the coiled-coil connector and the C terminal β- and γ-nodules, respectively. The scale bar represents 10 nm. **(D)** 100 superimposed SAXS *ab initio* bead models calculated by GASBOR under SAXS constraints. Beads randomly placed based on the X-ray scattering for the water hydration and protein are shown as red and black dots, respectively. The diameter of the C terminal β- and γ-nodules is about 90 Å. **(E)** Major classes of structure models generated based on SAXS and TM-HDX analysis are shown in cartoon representation.

In the 50-subclasses, fibrinogen micrograph averages consisted of a linear array of two distinct dimeric nodules at each C-terminus. Both nodules are held together with a thin connecting region, which approximately has a diameter of 20 to 30 Å (Fig. 3B, green and blue spheres and SFig. 5A). Both end nodules are comparable in size and herein represent the βC- and γC-nodules. The relative orientation of this doublet, with respect to the coiled-coil connector, is invariant in crystal structures (SFig. 6A). These distinct structural features of the C-terminal nodules observed in negative stain EM averages become fuzzier or disappear completely, when the number of averages is enforced from 50 to 20 and then to 10 (SFig. 5B and C). Approximately half of the population has a linear central dimeric interface about the central nodule (Fig. 3A and B). The most dominant averages i) are straight (Fig. 3B, 3^rd^ panel), ii) have a single kink either in the centre (Fig. 3B, 2^nd^ panel), iii) are in the close vicinity of a nodule at one end (Fig. 3B, 1^st^ panel), or iv) have two kinks (Fig. 3B, 4^th^ panel) on the fibrinogen scaffold. Some EM averages with two kinks have an oblique orientation, intermediate between the view with the end-nodule parallel to the grid or the view on the side (Fig. 3B, panels 4). Obliqueness was evident by the shorter length of the βC- and γC-dimer and the lack of two separate densities for each βC- and γC nodule (Fig. 3, panel 4) compared with a planar view (Fig. 3, panels 1-3). We speculate that in the presence of two flexions in fibrinogen, the βC- and γC nodules orient more out of the coiled-coil plane than in the presence of one or no flexion, giving rise to structurally heterogeneous conformers. Typically, sample heterogeneity is observed when a protein exists in different conformational states as exemplified by integrin receptors.[47, 48] Since there has been limited structural heterogeneity reported in fibrinogen crystal structures across different species,[3-5, 12] we next decided to further examine the overall fibrinogen’s conformation using solution X-ray scattering.

### X-ray Scattering of Fibrinogen Provides a Dynamic Conformational Ensemble Model

X-ray scattering profiles of the fractions from the peak of the SEC elution were collected and averaged (SFig. 7A and B, Table 2). The Kratky plot shows a bell-shape peak in the low-q region and does not converge to the q-axis showing that fibrinogen fractions used in our SAXS data collection correspond to a monomeric and multi-domain protein with pronounced domain-domain flexibility (SFig. 7C). D_max_ calculated from the short Guinier region is around 448 Å (SFig. 7D).

To gain further insight into the conformation(s) of fibrinogen in solution, we first generated *ab initio* models reconstructed from a chain-like ensemble of dummy residues without any secondary structure definition and generated 104 different SAXS models with explicit solvent and electron density maps using GASBOR.[49] Superimposed models revealed a Y-shaped, predominantly asymmetrical density distribution (Fig. 3D), and clustering *ab initio* models into groups containing similar volumetric distribution resulted in four major classes, which can be distinguished by the rotational pivot locations along the central axis. Effective radii (D_max_) of the *ab initio* models are 3 to 4 nm smaller than the previously reported fibrinogen length of 45 nm.[50]

In the central nodule dimer interface region, a prominent hydration shell was observed (Fig. 3D, red beads). When examined independently, the central axis adopted a highly flexible structure and the calculated dummy atom distributions, with two enlarged and globular nodules are similarly observed in EM class averages. From the resulting model, the central long-axis of each *ab initio* model has a width ranging ~30-40 Å, which is comparable to the width of the coiled coil-connector measured in our EM averages (Fig. 3A and B) and crystal structures. [3-5, 12] The average diameter of the combined C-terminal nodules is approximately 90 Å, which is also consistent with the size of the βC- and γC-nodule within a D region (Fig. 3D). From these models, we observed a highly labile globular region protruding from the central axis, which we assign to the structured portions of the globular αC-domain. This assignment is consistent with the TD-HDX classification of the αC domain (α:350-570), which is ascribed with rapid, temperature-independent exchange properties (class III behavior-Fig. 2C). Our secondary structure prediction and homology modelling suggest a domain-like character for residues in the range of 350-570 with β-helix propensity [51], which measures approximately 20-30 Å in diameter, consistent with the size and existence of these extra nodules.

### Architectural flexibility of fibrinogen

For flexible protein scaffolds like fibrinogen, modelling *in-solution* states as an ensemble of conformers represents a more holistic view of its conformational dynamism. Thus, to gain further insights into the *in-solution* dynamics of fibrinogen, we used a data-assisted assembly approach to demarcate the extent of flexibility under a set of constraints, which are derived from our HDX, EM and SAXS data. In generating rigid body models of fibrinogen heterodimer, linker (flexible) regions on the starting Fibrinogen homodimer model were defined as the following loops; A1-V39, R95-N103, L204-D213, M235-W315 for the α-chain, Q1-G64, N137-E141 and V199-E210 for the β-chain, and Y1-E13, N69-P76 and D141-D152 for the γ-chain. The linker regions connected 20 different monomeric rigid modules. Using the simulated annealing program, CORAL [49], translation and rotation of these rigid modules were carried out to obtain structure models with optimal positions and orientations of the fibrinogen homodimer scaffold that conform to the restraints from experimental X-ray scattering. The resulting structure models exhibited well-converged **χ-**values (SFig. 7E) and appear asymmetrical. Structure models showed one or two hinge regions along the axis at the coiled-coil connectors and/or, to a lesser extent, about the central nodule of the E-region, and clustered into four major conformationally related classes (Fig. 3E).

### Visual insights into the fibrinogen models through low-resolution data comparison

We quantitatively examined the correlation of i. EM averages with the structure models that were generated under SAXS and TM-HDX constraints (Fig. 3E compared to 3B) and ii. structure models with the *ab initio* models that were generated under only SAXS constraints (Fig. 3D compared to 3E). To compare structure models with EM averages, first filtered molecular envelopes of the class representatives of structure models at 20 Å resolution were used to calculate regularly spaced pseudo-EM projection at 2° intervals and then cross-correlated with the EM averages. To compare the *ab initio* models with these structure models, the *ab initio* bead models were converted to volumetric situs file format and cross-correlated to the structure models using Corales. [52]

The EM class averages, clusters of structure and *ab initio* models agree well, with four overall topological classes identified for fibrinogen; *i*. straight, *ii*. central-bent, *iii*. straight-bent and *iv*. trans homodimers (Fig. 4A-D, cartoons). The dimer interface of fibrinogen in most EM averages is largely linear (Fig. 4 A and B-panels 1 and 2), consistent with structure and *ab initio* models (Fig. 4 A and B-panels 3-5), though 35% of EM averages, and some structure, and *ab initio* models showed bending at the dimer interface (Fig. 4C, panels 1-5). The majority of EM averages (75%) showed pronounced bending along the coiled-coil axis (SFig. 5A), and cross-correlated well with similar SAXS class-representatives displaying coiled-coil bends over other possible conformers (Fig. 4A, B and D). The trans-topology was also detected, albeit rarely, in EM images (Fig. 4D, panels 1 and 2) and structure models (Fig. 4D, panels 3 and 4) and *ab initio* models (Fig. 4D, panel 5) with the coiled-coil bent conformations.

**Figure 4.**
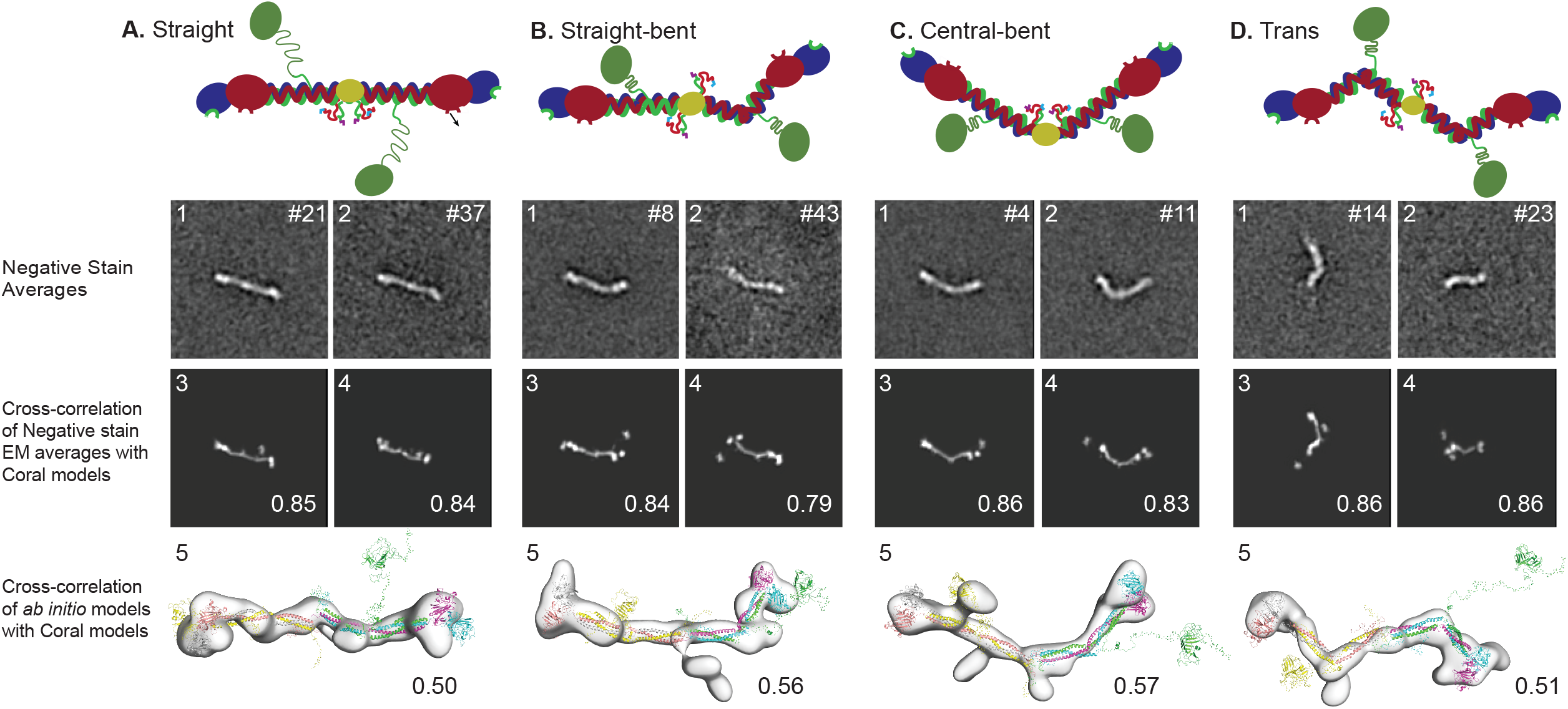
Comparison of EM averages, structure and *ab initio* fibrinogen models. Four major classes detected adopt **(A)** straight, **(B)** straight, bent **(C)** central bent and **(D)** trans configurations. Cross-correlation coefficients between negative stain EM averages and structure models, and between structure models and *ab initio* models are noted left corners of each panel.

Next, we calculated angle distributions between the defined globular modules to quantitatively describe their relative angle deviations along the central axis of the fibrinogen homodimer scaffold. We constructed histograms that present the angle distributions and fits between *i*. the coiled-coil-E domains, extending from both sides of the centre of the fibrinogen molecule, *ii*. the coiled-coil-E domain and the coiled-coil D domain, *iii*. the coiled-coil D domain and the dimeric βC- and γC-nodules and *iv*. the coiled-coil D domain and the αC-domain (Fig. 5). In defining an angle between two rigid bodies, the centre of mass (COM) for the specified domains were initially determined using PyMOL and the angles were calculated between two COMs and the connector residue(s) between two domains. The dimer interface in the central nodule of fibrinogen, where five disulphide bridges connect the N-termini of the six chains, adopts two different topological conformers, one (*major conformation*) with a nearly straight angle (178° ± 14°) and the other (*minor conformation*) with an obtuse angle (132° ± 11°) (Fig. 5A). All resolved crystal structures adopt a straight homodimerization interface,[3-5, 12] consistent with the major conformation reported in our structure models. Flexibility was evident along the coiled-coil connector, resulting in three major conformers that adopt angles between the coiled-coil-E domain and the coiled-coil-D domain (177° ± 22°, 94° ± 12° and 130° ± 14°) (Fig. 5B). The relative orientation between the coiled-coil-connector and the βC- and γC-nodule dimer appears to adopt a single mode with an angle of 129° ± 26° (Fig. 5C). Also, a broad and bimodal histogram was observed for the angle defined between the αC-domain and the coiled-coil connector (180° ± 30° and 101° ± 31°) (Fig. 5D). In sum, there are major flexion points, located about the centre of the fibrinogen molecule and in the centre of the coiled-coil connector, consistent with the TD-HDX identified flexion. In all models, the αC-domain, as expected, appears to move almost freely and independently. The latter property provides a likely explanation for why the αC-domain is typically averaged out in our EM averages (Fig. 3B, SFig5A-C), though, select raw, unfiltered images display faint density protruding from the coiled-coil axis (Fig. 3A).

**Figure 5.**
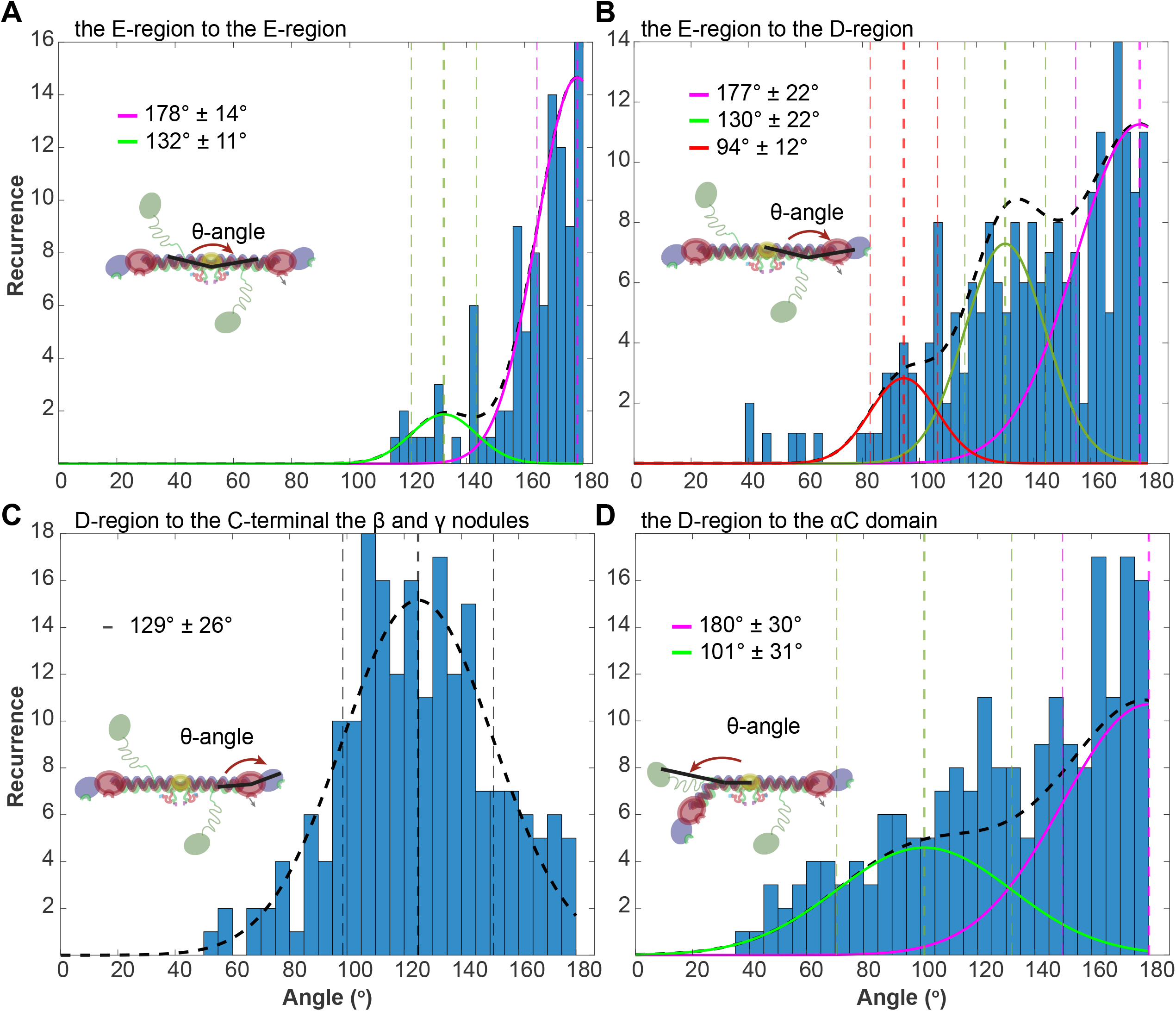
Distributions of the fibrinogen bending angles. Angles between the center of mass (COM) for the specified domains were calculated using PyMOL. Histogram fits for each θ-angle between the **(A)** E-region/E-region, **(B)** E-region/D-region, **(C)** D-region/C-terminal the β and γ nodules and **(D)** D-region/αC-domain were performed using Matlab, and mean and st.dev of fits are noted above the cartoon showing the bending by a curved arrow.

## Discussion

Structural information of fibrinogen gleaned from over seven decades and from different species using a variety of techniques has culminated with the consensus that its structure is mostly linear, with limited flexibility along the coiled-coil regions (a “planar sigmoidal” shape is commonly used to describe the slight bend/twist in this region). In addition, the relative positions of the C-terminal β- and γ-nodules to each other and orientation of the βC- and γC-nodules with respect to the coiled-coil regions are largely invariant. In contrast, the studies of fibrinogen self-diffusion combined with MD simulations showed unusual and heterogenic hydrodynamic behaviour with a smaller radius.[19] Fibrinogen structural heterogeneity has been also suggested by previous adsorption-based studies using AFM, transmission electron microscopy (TEM), and MD simulations.[20, 42] Our independent experimental testing of the full range of fibrinogen dynamics for an intact, solution sample provides novel structural and mechanistic insight into the “*Fibrinogen dynamism*” and has pinpointed two significant flexions: one located at the coiled-coil connector at the intersection of the D- and E-regions and the other located at the homodimerization interface that connects two E-regions of the individual fibrinogen monomers. These multiple pivoting flexions are most likely the underlying reason for the observed reduction in the fibrinogen radius by 3-4 nm in solution as probed herein and by Zuev et al., [19] in comparison to the crystal and EM structures wherein fibrinogen is in a crystalline lattice or immobilized to grids.

Solution phase hinging about the centre of the fibrinogen molecule has been suggested by previous AFM and TEM studies, [18, 53] but whether these were artifacts of the fibrinogen-surface interaction intrinsic to these techniques was unclear; crystallographic structures of human fibrinogen are linear. Our solution-phase SAXS data, along with our EM structures, clearly show that while most molecules adopt a relatively straight configuration, a small population are bent about the central nodule (i.e., the dimer interface), at an angle of ~130°. Flexibility within the coiled-coil connector was dramatic. The planar sigmoidal shape in various fibrinogen crystal structures revealed an angular range of 176° ± 1.3° for the coiled-coil connector region, which agrees well with one of our observed populations (177° ± 22°). However, we resolved to other orientations showing dramatically more bend in this region with angles of 94° ± 12° and 130° ± 14°; these conformations are in agreement with the TD-HDX data and orientations previously predicted by molecular dynamics simulations, but have not been previously reported on in experiments (Fig. 5B). [20] Secondly, hinging at the coiled-coil region leads to rotation of the βC- and γC-nodule on the axis of the coiled-coil, permitting an orientation that exposes unique, and previously undetected, conformations. The extent of the hinging, both rotational and translational, at the fibrinogen flexions likely assists and uniquely establishes alignment of the knobs and holes during the double-stranded protofibril formation—the initial step in the formation of the fibrin network—and thus, contributes to and regulates the overall flexibility of protofibrils that has been observed in AFM [54, 55] and EM images [56–58].

Two N-glycosylation sites, γ-N52 and β-N364, have been identified in human fibrinogen, with the former located proximal to the coiled-coil flexion and the latter being flanked by peptides with class II and III (intermediate and fast) hydrogen exchange behaviours. Also, two unique binding motifs, α-^95^RGD^97^ and α-^17^GPR^19^, are located in the coiled-coil flexion next to the N-glycosylation site, γ-N52 and the flexion positioned in the homodimerization interface, respectively. Any modification of these sites (e.g., post-translational modifications or point mutations) appears to alter the bulk functional properties of fibrinogen in solution, as exemplified by the N-glycans modifications leading to aforementioned diseases, α-^95^RGD^97^ mutations in promoting only fibrinogen-mediated adhesion and retraction of platelets but not platelet aggregation, and α-^17^GPR^19^ mutations in the myeloid leukocyte activation. {Loike, 1991 #796} Apart from αXβ2 and αIIbβ3 integrins, fibrinogen interacts with an array of other integrins and proteins. The integrin-specific cryptic sides are located in i. the αC region interacting with α5β1[21], αVβ3[22], αVβ5[59] and αIIbβ3, ii. the D-region interacting with αXβ2 and iii. the γ-nodule interacting with αMβ2. However, the molecular basis of these fibrinogen interactions remains elusive due, in part, to the cryptic nature of the sites on fibrinogen. For example, the αDβ2 binding region has yet to be determined. These observations in the context of the fibrinogen internal conformational flexibility suggest that fibrinogen-structural transitions between different states via “conformational breathing” is essential in regulating the accessibility of receptor binding epitopes, clot architecture and growth kinetics. This breathing would also help to facilitate other structural transitions such as the hypothesized “pull out” of the αMβ2 binding site from the γ-nodule.[29]

The βC- and γC-nodules are structural homologs (all-atom RMSD for the superimposed βC- and γC-nodules is 0.86 Å, PDB# 3ghg) and have a rigid interface between each other and with the coiled-coil connector in all crystal structures.[3–5, 12] A single geometric position of the βC- and γC-nodules are also clear in our EM images. Given that the tilt angle between the coiled-coil-connector and the βC- and γC-nodule dimer is ~135° in crystal structures and the rotational swap of the βC- and γC-nodules during the simulated annealing with SAXS restraints resulted in a mean angle deviation of 129°, the relative orientation between the coiled-coil-connector and the βC- and γC-nodule dimer in both solution and immobilized states appears rigid. This local rigidity most likely conformationally shields or shortens the temporal exposure of the plasminogen/tPA site α148-160 in fibrinogen from premature cleavage by plasminogen. Since plasminogen can readily access this site in the insoluble fibrin state, the rigidity of the interface formed between the dimeric βC- and γC-nodules with the coiled-coil connector also appears to be functionally important, although the broad distribution of angles (Fig. 5C) accessible to this region suggests that it could “breath” to some extent as well.

Despite a vast experimental and clinical literature revealing the epidemiology and genetics of congenital fibrinogen diseases, which define the fibrinogen-mediated disorders, a better understanding of *in-solution* conformational ensemble and dynamisms of fibrinogen is expected to shed important insight into the significant clinical heterogeneity in the severity of the thrombotic and hemorrhagic pathologies. Three types of pathologies stem from fibrinogen mutations: dysfibrinogenemia, hypofibrogenemia and afibrinogenemia. The phenotypic symptom severity spectrum of these diseases in clinics, such as in patients with dysfibrinogenemia is broad; some patients have severe bleeding episodes, and thrombotic phenotypes in others are observed, whilst some stay asymptomatic during their whole lifespan (Fig. 6B). Interestingly, phenotypic severity of these fibrinogenemia may be linked to the location(s) of the point mutations; most of the well-tolerated mutations, showing asymptomatic to mild dysfibrinogenemia, are positioned adjacent to the flexion regions. For instance, α-R95 to E or Q mutations showed a slight reduction in the α-helix content in comparison with wild-type (WT) fibrinogen, [60] and α-R95K mutation resulted in asymptomatic to mild dysfibrinogenemia. [61]. On the other hand, β-R166C and β-L172Q mutations that are positioned in the highly rigid molecular structures, found in the Class I region of our HDX map —similar rigidity identified for the plasminogen/tPA proteolysis site—are attributed to impaired polymerization, leading to severe and abnormal clotting. In sum, fibrinogen, due to its intrinsic conformational flexibility, likely exists in a dynamic equilibrium accessing four major conformations (Fig. 4), with mutations and PTMs regulating this equilibrium and molecular interactions of fibrinogen temporally and spatially.

**Figure 6.**
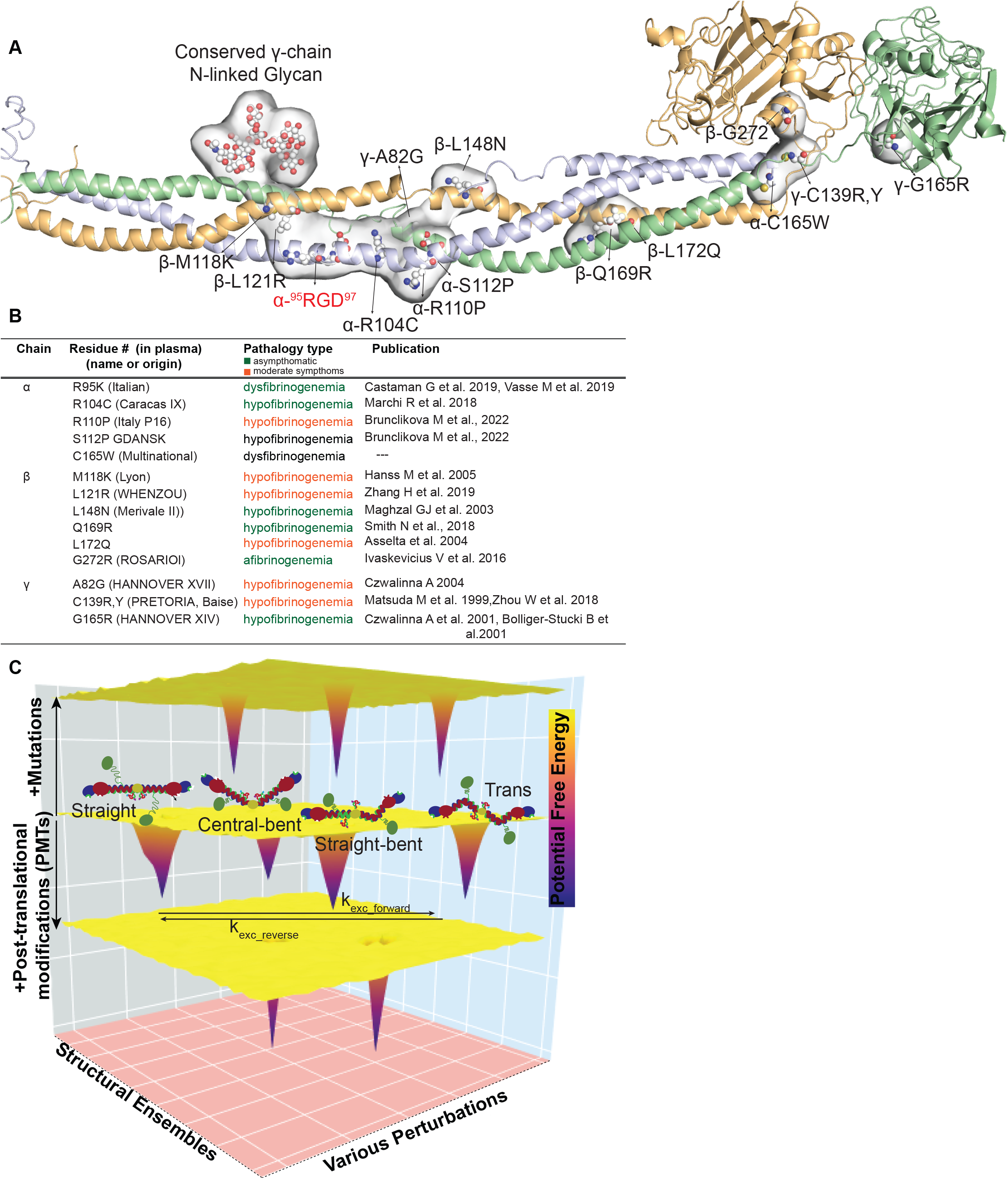
Single mutations across the phenotypic severity of the congenital fibrogenemia and conformational energy landscape of fibrinogen. **(A)** Fibrinogen structure showing residues of the modelled γN52 N-linked glycan, α-^95^RGD^97^ cryptic binding epitopes, and single/homozygous mutations that implicated only in mild or asymptomatic Fibrinogen-related pathologies as CPK representation on the cartoon backbone. **(B)** Name, pathological severity and reference of these mutations are noted. **(C)** The intrinsic conformational flexibility of fibrinogen, as observed in four major conformers, is depicted on the hypothetical potential energy landscape.

It is most interesting that the flexion in the coiled-coil region also appears highly diverged across species, likely acting in analogy to a mutational hotspot and providing a unique environmental, physical, and/or metabolic adaptation to each species. For instance, avian species (birds) lack the α-^95^RGD^97^ motif in this specific flexion. Since birds can acutely rise and chronically maintain their blood pressure during flight by barodenervation using aortic baroreceptors, and their thrombocytes (nucleated blood cells that function like platelets) are much larger, [62] the lack or loss of the α-^95^RGD^97^ motif appears to be advantageous in birds’ cardiovascular system, preventing immediate clotting[63] and tight aggregate formation,[64] and thus, providing a better (slower) control of bleeding.

Based on our in-solution studies, fibrinogen has shown a breadth and depth of structural diversity even at the low spatial resolution, and the newly identified flexions are populated with cryptic binding epitopes, PTMs, and well-tolerated mutations. Accordingly, our hypothesis is that fibrinogen allosteric mechanisms are constructed by the properties of the native potential energy landscape of the system, and the extent of distortion of this landscape via various types of perturbations alters the fibrinogen functional outputs (e.g., change in the clotting architecture, temporal accessibility of epitopes and pathologic severity of phenotypic manifestation) (Fig. 6C). Major components that alter the fibrinogen potential energy landscape are hence i. relative Gibbs free energy of fibrinogen conformers that consists of cryptic-ligand binding competent and incompetent conformations, ii. transition kinetics between these conformers, and iii. the intrinsic affinities of interacting molecules, which could also possibly remodel the fibrinogen energy landscape. Therefore, studies that link the roles of fibrinogen conformational dynamism and kinetics, timescale, and energy barriers among fibrinogen-conformers to the functional outputs in health and pathologies would be essential in better understanding fibrinogen biology and developing more effective fibrinogen therapeutics that fine-tune fibrinogen functions in various haematological disorders.

## Methods

### Isolation and characterization of human Fibrinogen peak 1

We studied the full-length human fibrinogen molecule containing the α, β and γ-chains spanning from Ala1-P625, Q1-Q461, and Y1-V411, respectively. This protein was sub-fractionated, and the major 1^st^ peak, containing two platelet-binding ‘γ-A’ γ-chain isoforms, is free from any fibrinogen-binding molecules, such as factor XIII (Enzyme Research Labs, South Bend, IN). Prior to each of the structural methods used in the following sections, this fibrinogen sample was further purified to homogeneity by size-exclusion chromatography (SEC). The resulting fibrinogen yielded a single, dimeric (AαBβγ)_2_ peak and in reducing SDS-PAGE, this SEC fraction showed three bands corresponding only to the α, β and γ-chains (SFig. 3 A, B). The CD spectrum of this fibrinogen peak sample (not shown) exhibits a well-defined fold indicative of predominantly alpha helices, consistent with the structural model. The sample is also functional; the clottability of peak 1 fibrinogen was found to be ≥ 90%. Clottability is a standardized measure for the functional integrity of fibrinogen and is determined spectrophotometrically by the loss of soluble fibrinogen sample to insoluble fibrin through the treatment with thrombin.

### HDX sample preparation

Aliquots of fibrinogen, frozen after the FPLC purification, were thawed and were diluted 10-fold in 10 mM HEPES, 0.5 mM EDTA, pD 7.4 D_2_O (99%D, Cambridge Isotopes) buffer (corrected; pD = pH_read_ + 0.4). Samples were incubated at a specified temperature (temperatures studied: 10 and 25°C), using a water bath, for 10 time points (0, 10, 20, 45, 60, 180, 600, 1800, 3600, and 7200 s) at each temperature. For each temperature, the time points were collected over the course of three days and randomized to reduce systematic error. Each sample (from a unique time point) for a given temperature was prepared and processed once.

Upon completion of the designated incubation time, all samples were then treated identically; the samples were rapidly cooled (−20°C bath) and acid quenched (to pH 2.4, confirmed with pH electrode, with 0.32 M citric acid stock solution [90 mM final concentration] on ice). Procedures from this point were conducted near 4°C. Prior to pepsin digestion, the samples were incubated with guanidine HCl (in citric acid, pH 2.4) and TCEP to a final concentration of ca. 0.5 M and 1 M, respectively, for 5 minutes. The addition of the reducing and chaotropic agents was necessary for obtaining peptide coverage of ≥70% per chain. Fibrinogen samples were digested for 2.5 min with immobilized pepsin that had been pre-equilibrated in 10 mM citrate buffer, Ph 2.4. The peptide fragments were filtered, removing the pepsin, using spin filters with cellulose acetate membranes and centrifugation for 10 seconds at 4°C. Samples were flash frozen immediately in liquid nitrogen and stored at −80°C until data collection.

### Liquid chromatography-tandem mass spectrometry for peptide identification

To identify peptide fragments of fibrinogen resulting from pepsin digestion, samples of pepsin-digested fibrinogen at time = 0s (H_2_O buffer) were analyzed using a Thermo Dionex UltiMate3000 RSLCnano liquid chromatograph (LC) that was connected in-line with an LTQ Orbitrap XL mass spectrometer equipped with an electrospray ionization (ESI) source (Thermo Fisher Scientific, Waltham, MA). The LC was equipped with a C18 analytical column (Acclaim® PepMap 100, 150 mm length × 0.075 mm inner diameter, 3 μm particles, Thermo). Solvent A was 99.9% water/0.1% formic acid and solvent B was 99.9% acetonitrile/0.1% formic acid (v/v). The elution program consisted of isocratic flow at 2% B for 4 min, a linear gradient to 30% B over 38 min, isocratic flow at 95% B for 6 min, and isocratic flow at 2% B for 12 min, at a flow rate of 300 nL/min. The column exit was connected to the ESI source of the mass spectrometer using polyimide-coated, fused-silica tubing (20 μm inner diameter × 280 μm outer diameter, Thermo). Full scan mass spectra were acquired in the positive ion mode over the range *m/z* = 350 to 1800 using the Orbitrap mass analyzer, in profile format, with a mass resolution setting of 60,000 (at *m/z* = 400, measured at full width at half-maximum peak height).

In the data-dependent mode, the eight most intense ions exceeding an intensity threshold of 30,000 counts were selected from each full-scan mass spectrum for tandem mass spectrometry (MS/MS) analysis using collision-induced dissociation (CID). Real-time dynamic exclusion was enabled to preclude the re-selection of previously analyzed precursor ions. Data acquisition was controlled using Xcalibur software (version 2.0.7, Thermo). Raw data were searched against the amino acid sequence of fibrinogen α/β/γ chains using Proteome Discoverer software (version 1.3, SEQUEST, Thermo) to identify peptides from MS/MS spectra.

### Liquid chromatography-mass spectrometry for hydrogen/deuterium exchange measurements

Deuterated, pepsin-digested samples of fibrinogen (see details above) were analyzed using an Agilent 1200 LC (Santa Clara, CA) that was connected in-line with the LTQ Orbitrap XL mass spectrometer (Thermo). The LC was equipped with a reversed-phase analytical column (Viva C8, 30 mm length × 1.0 mm inner diameter, 5 μm particles, Restek, Bellefonte, PA) and guard pre-column (C8, Restek). Solvent A was 99.9% water/0.1% formic acid and solvent B was 99.9% acetonitrile/0.1% formic acid (v/v). Each sample was thawed immediately prior to injection onto the column. The elution program consisted of a linear gradient from 5% to 10% B over 1 min, a linear gradient to 40% B over 5 min, a linear gradient to 100% B over 4 min, isocratic conditions at 100% B for 3 min, a linear gradient to 5% B over 0.5 min, and isocratic conditions at 5% B for 5.5 min, at a flow rate of 300 μL/min. The column compartment was maintained at 4 °C and lined with towels to absorb atmospheric moisture condensation. The column exit was connected to the ESI source of the mass spectrometer using PEEK tubing (0.005” inner diameter × 1/16” outer diameter, Agilent). Mass spectra were acquired in the positive ion mode over the range m/z = 350 to 1800 using the Orbitrap mass analyzer, in profile format, with a mass resolution setting of 100,000 (at m/z = 400). Data acquisition was controlled using Xcalibur software (version 2.0.7, Thermo).

Mass spectral data acquired for HDX measurements were analyzed using the software, HDX WorkBench. [65] The percent deuterium incorporation was calculated for each of these peptides, taking into account the number of amide linkages (excluding proline residues) and the calculated number of deuterons incorporated (see SI HDX Table). The values were normalized for 100% D_2_O and corrected for peptide-specific back-exchange as reviewed in. [66] The data were plotted as deuterium exchange versus time using Igor Pro software. HDX traces are available in the SI HDX traces.pdf

### Synchrotron SAXS Measurements

Purified fibrinogen purchased from the Enzyme Research Laboratories were shipped on ice to the synchrotron and incubated at ambient temperature for 15-20 min before x-ray solution scattering measurements were performed. For the SEC-SAXS experiment, 200 μL of 10 mg/mL fibrinogen was injected into the Superdex S200 column for each experimental run. The mobile phase was 20 mM HEPES, 150 mM NaCl, and pH 7.4. The SAXS experiments on the prepared samples were collected using National Synchrotron Light Source-II (NSLS-II) Beamline 16-ID (LiX) at Brookhaven National Laboratory. Despite the short Guinier region, the Kratky plot of fibrinogen showed a folded multidomain protein with intrinsic flexibility (Fig. 4D). *I*(0) and the pair distance distribution function *P*(*r*) were calculated by circular averaging of the scattering intensities *I*(*q*) and scaling using the software GNOM [49]. Fibrinogen scattering data was processed to a q(A^-1^) of 0.09 and 1.15.

### Ab inito and structural modelling of the Fibrinogen conformations under SAXS constraints

*Ab initio* modelling software, GASBOR software from the ATSAS package were used to generate 100 bead models. Those models then were then aligned using DAMAVER or DAMCLUST. [49]

For rigid body modelling, Complexes with Random Loops (CORAL) were used to create rigid body conformers with flexible or hinge regions conforming with SAXS data. These hinge regions that are interdomains in a protein architecture and help define freely moving rigid bodies were determined based on our TD-HDX and sequence divergences, ultimately defining 20 different rigid bodies. These rigid bodies for CORAL models were defined as follows; For Aα chain: V20-L94, R104-P203, V214-S569, and N316-S569. For the β chain, the following residues were paired as rigid bodies: C65-E136, Y142-T198, and C211-F458. As for the γ-chain residues:

R1-D58, N64-K127, and C140-I381 were selected to act as rigid bodies. In crystal structures, the βC- and γC-nodules always adopt the same relative configuration to each other, hence the βC- and γC-nodules were kept as stable-dimer. Additionally, both E- and D-region sections of the coiled-coil connectors also have rigid quaternary scaffolds, so three single-bond constraints between α-β, β-γ, and γ-α chains for the E and D region sections of the coiled-coil connectors were introduced. Via simulated annealing, the most optimal positions, and orientations of high-resolution fibrinogen molecules were obtained with the input parameters for the modelled rigid domains and the missing peptide portions of the flexible linker residues, and more than one hundred conformers were generated.

### Negative stain EM of fibrinogen

Within 2 hours after Superdex 200 10/30 column purification of ~30 μg of fibrinogen in 20 mM Hepes pH 7.4 and 150 mM NaCl were adsorbed to glow discharged carbon-coated copper grids, stained with uranyl formate, and inspected with a Jeol2100 operated at 200 kV. Low-dose images were acquired with a JEOL2100 microscope at 200 kV and a nominal magnification of 12,000X or 52,000X using a defocus of –1.5 μm. About 3,000 particles were interactively picked, windowed into individual images using the BOXER module of EMAN [67], and subjected to 10 cycles of multi-reference alignment and *K*-means classification into 50, 20 or 10 classes using SPIDER [46] as described. [48]

### 2D and 3D cross-correlations of rigid body models with EM averages and ab inito models

For 2D cross-correlation of rigid body models with EM averages, filtered molecular envelopes at a resolution of 20 Å from representative rigid body model structures from the earlier cluster analysis of the SAXS-based rigid body models were used to back-calculate regularly spaced negative EM projections at 2^°^ oscillations and were cross-correlated with EM class averages using SPIDER. [46]

For 3D cross-correlation of rigid body models with *ab initio* models, bead models were converted to the situs format that represents the *ab initio* models as volumetric structures. 3D Cross-correlation between *ab initio* volumes and rigid body models was calculated using the Exhaustive One-At-A-Time 6D search tool, colores. [52]

### Angle Measurements between rigid bodies

Angles between the rigid bodes/domains were structure models generated by SAXS-based rigid body models using PyMOL script *Angle_between_domains*. [68] Rigid bodies defined are 1) the coiled-coil region in the E-region, 2) the coiled-coil region in the D-domain, 3) β- and γ-nodules and 4) the αC-domain. Angles measured are between the rigid bodies of 1 and 2, 2 and 3, 1 and 4. Lastly, we also measured the angle at the homodimerization interface (1 and 1). Histogram plots showing these angles were fitted using MATLAB, using full and half-Gaussian and means and standard deviations were calculated for each angle distribution.

## Supporting information

SFig.

SI HDX Table

SI HDX traces

## Author Contributions

M.S., N.E.H. and A.R.O. conceived the work and wrote the manuscript. P.M. prepared the fibrinogen samples and help with EM and SAXS data acquisition. J.E.E.P. analysed the EM and SAXS data, wrote the manuscript. G.P. purified fibrinogen for HDX and SAXS data collection. K.R. assisted in HDX-MS analysis. A.T.I. collected MS data. Q.N. constructed histogram plots and full and half Gaussian fits.

## Acknowledgments

This work was supported by NIH grant NHLBI R15HL150666-01 to PI. A.R.O. and Co-Is, M.S. and N.E.H., NIH grant 1S10 OD020062 to A.T.I. and the National University Research Fund to M.S. We thank Danny Le, Mehmet Furkan Tasdelen, Jose Maria De La Garza, Nicholas Carter Kirby for their critical insights and technical help, and Baylor College of Medicine EM-core with negative stain EM data acquisition. This work was completed in part with resources provided by the Research Computing Data Core at the University of Houston. We also thank the NSLS-II beamline 16-ID (LiX) staff at the Brookhaven National Lab. The LiX beamline is part of the Center for BioMolecular Structure (CBMS) and is supported primarily by the National Institutes of Health (NIH), National Institute of General Medical Sciences (NIGMS) through a P30 grant (P30GM133893) and by the Department of Energy Office of Biological and Environmental Research (KP1605010). LiX also received additional support from NIH grant S10 OD012331. As part of NSLS-II, a national user facility at Brookhaven National Laboratory, work performed at the CBMS is supported in part by the US Department of Energy, Office of Science, Office of Basic Energy Sciences Program under contract number DE-SC0012704.

