## Supplementary material for "Chorography and conformational dynamism of the Soluble Human Fibrinogen in solution": SFig.

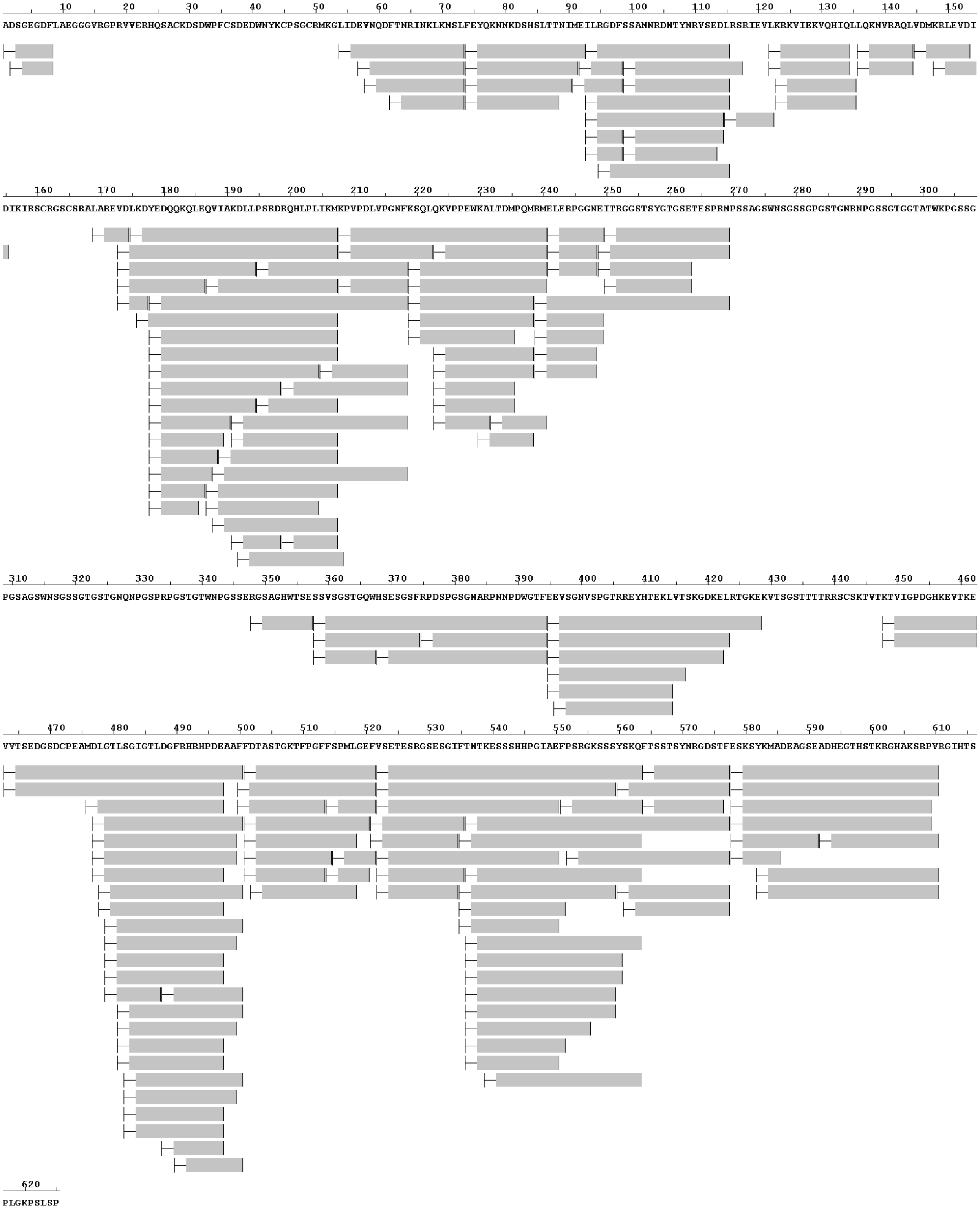


**Supplementary Figure 1. Complete peptide coverage map for fgn α-chain**

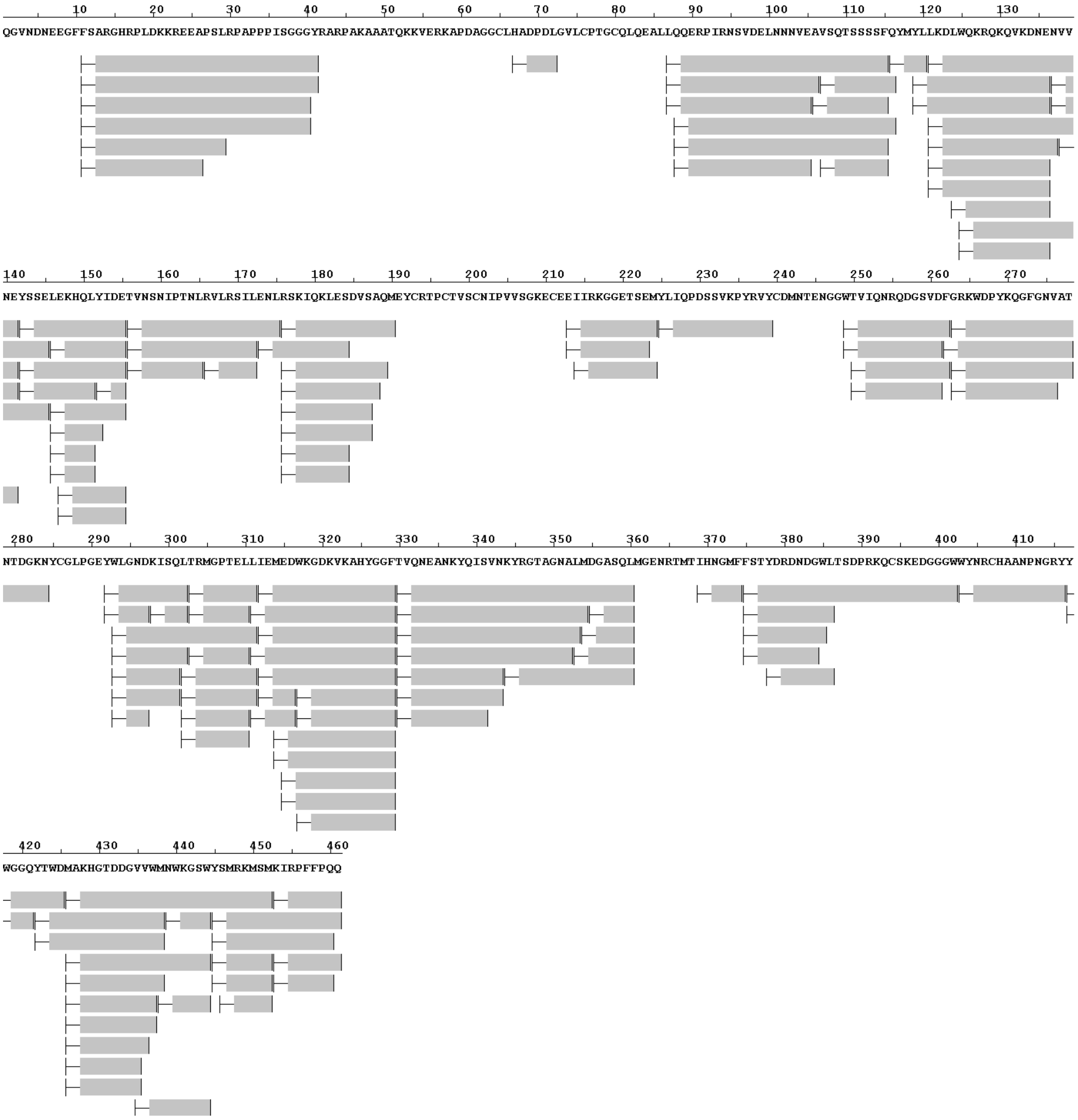


**Supplementary Figure 2. Complete peptide coverage map for fgn β-chain**

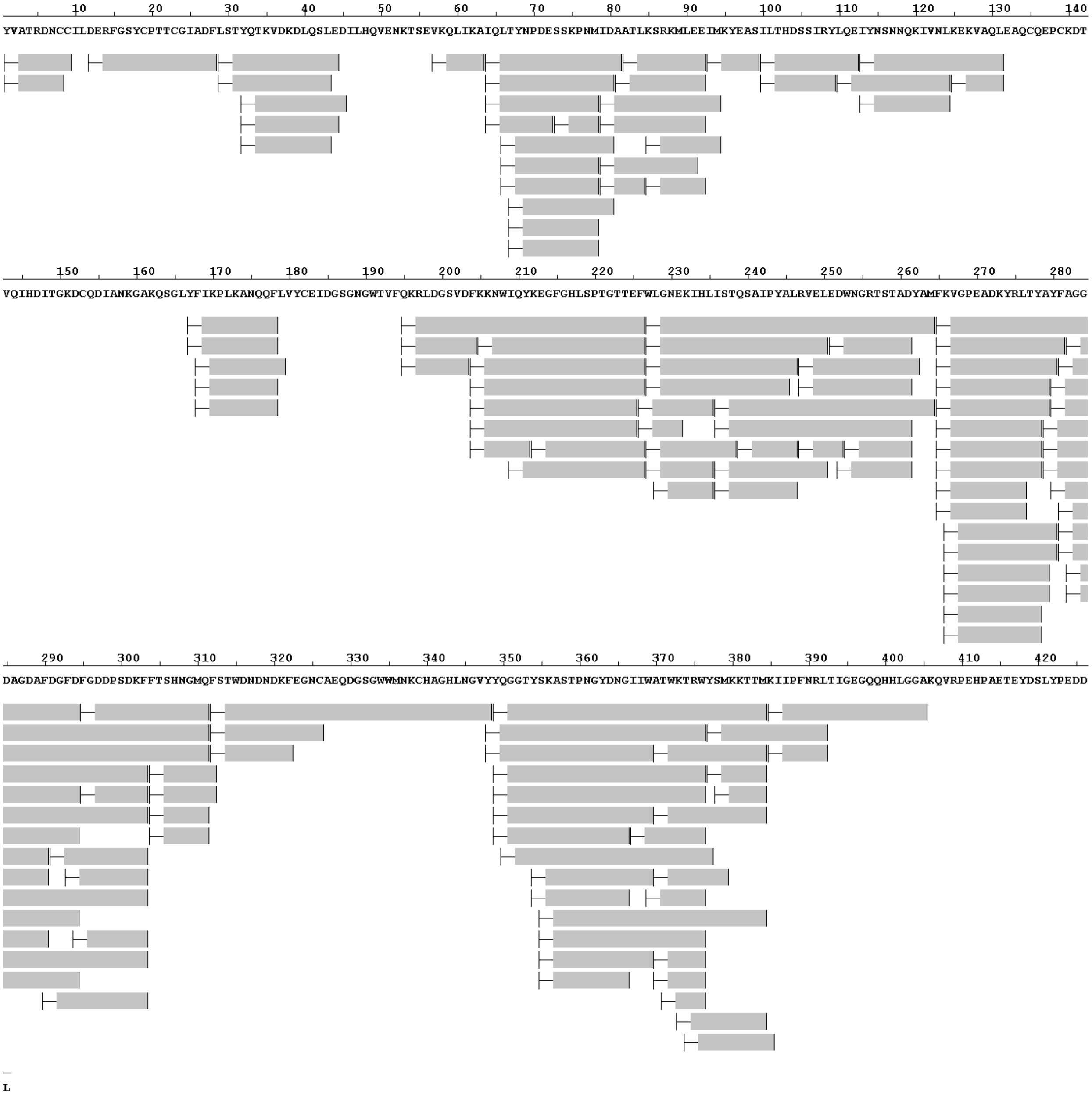


**Supplementary Figure 3. Complete peptide coverage map for fgn γ-chain**

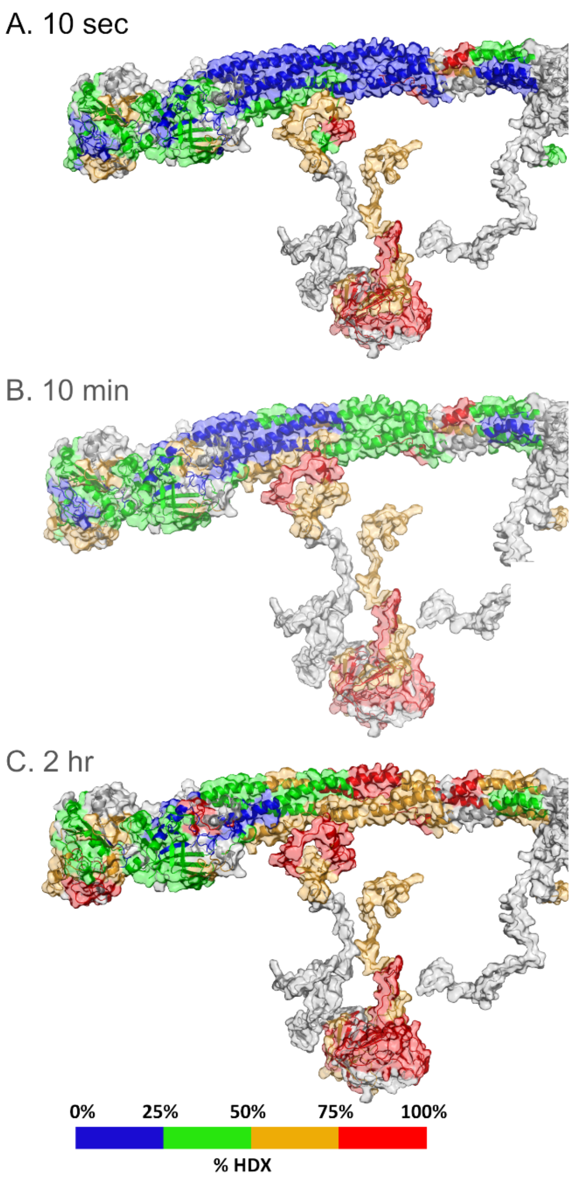


**Supplementary Figure 4. HDX-MS properties of Fgn at a single temperature**. Time evolution of the extent of HDX-MS of human fibrinogen, collected at 25 °C. The structural model of Fgn is colored according to the color bar. Peptides colored in gray are uncovered/unassigned in the mass spectra. Refer to the SI HDX file for the complete set of HDX-MS plots.

A

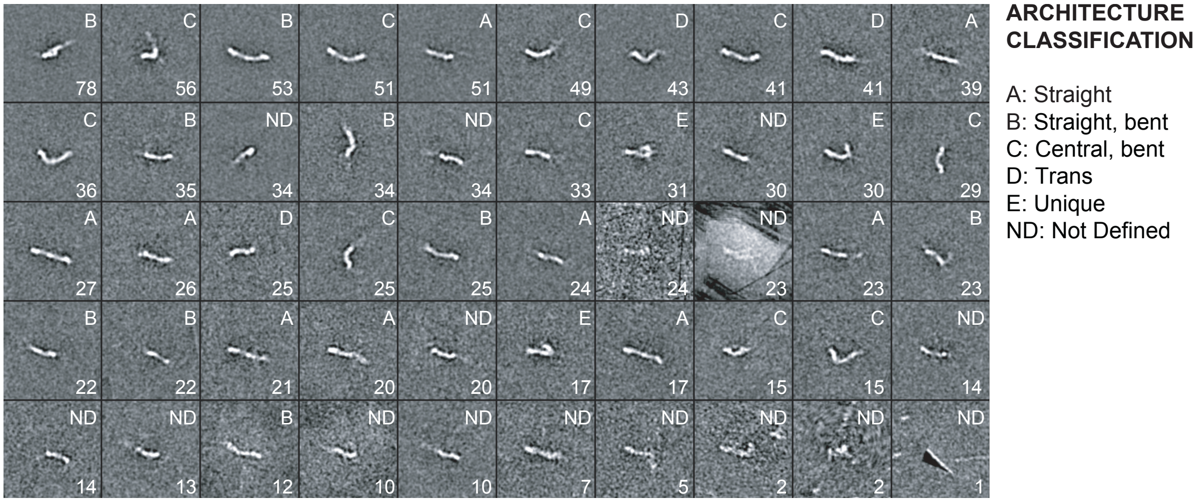


| Straight | Straight, bent | Central, bent | Trans |
|---|---|---|---|
| 248 (25%) | 304 (30%) | 350 (35%) | 109 (10%) |

B

C

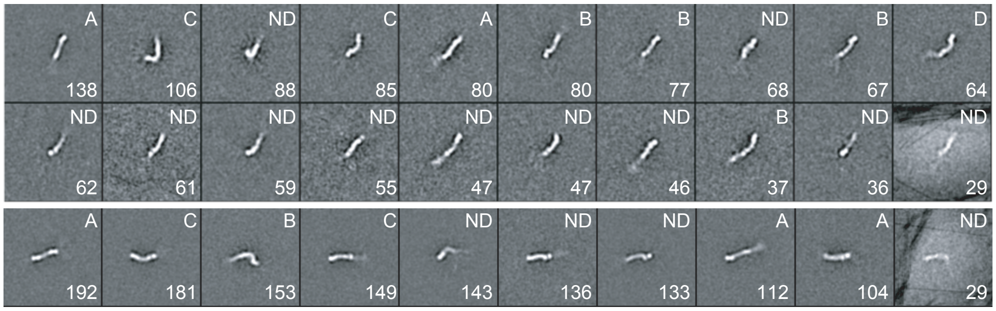


**Supplementary Figure 5.** Multi-reference aligned EM class averages of negatively stained fibrinogen. Total **(A)** 50, **(B)** 20, and **(C)** 10 EM class averages were generated, and classified based on their overall shapes. Number of total images for each major classes used during the multireference alignment were noted for only 50-class.

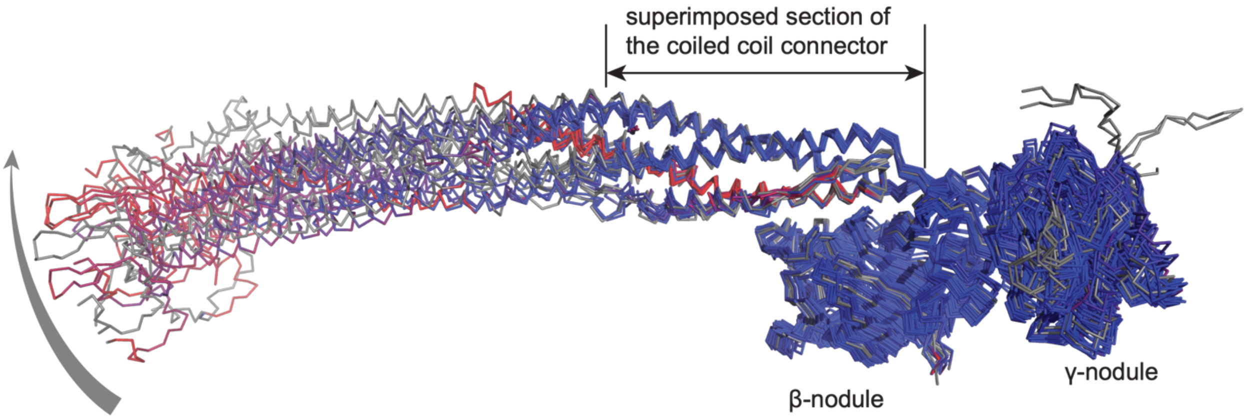


**Figure 6. Overlays of fgn X-ray structures**. Fibrinogen structures (1m1j, 1ei3 from chicken, 1deq from bovine, 3ghg from human) were superimposed on the D-region, colored by RMSD. While the E-region relative to the D-region shows high flexibility, the C-terminal nodules and their orientation relative to the D-region appears highly rigid.

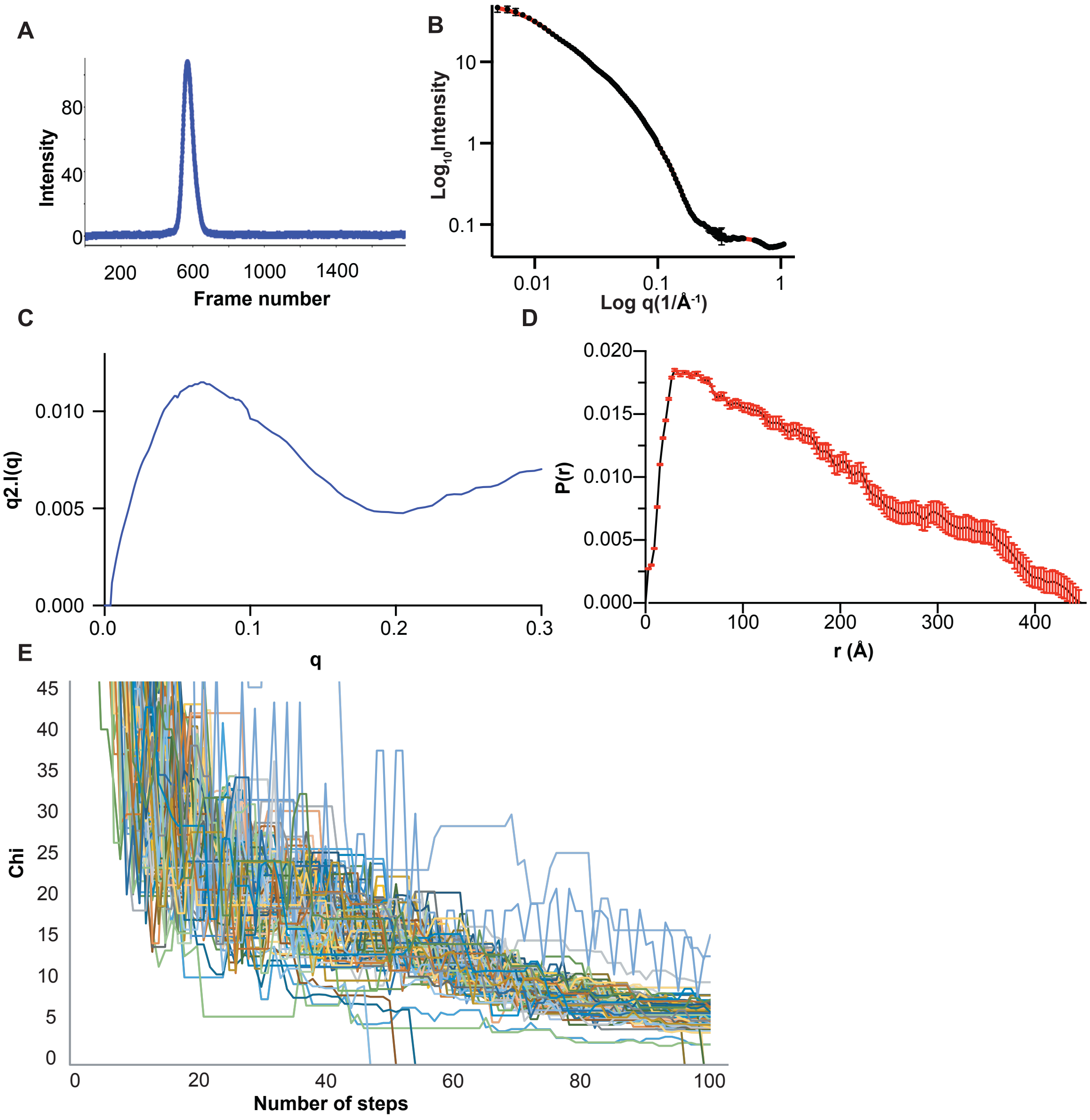


**Supplemenary Figure 7. SAXS data reduction and convergence χ-values of the structure models during the simulated annealing.**

**(A)** Size exclusion profile during the X-ray scattering data collection, **(B)** Raw intensity of I(q) SAXS data, **(C)** the pairwise distribution curves (P(*r*)) derived from the SWAXS and **(D)** Kratky plot. **(E)** The χ-value of each structure model during each 100 steps of the simulated annealing were plotted.

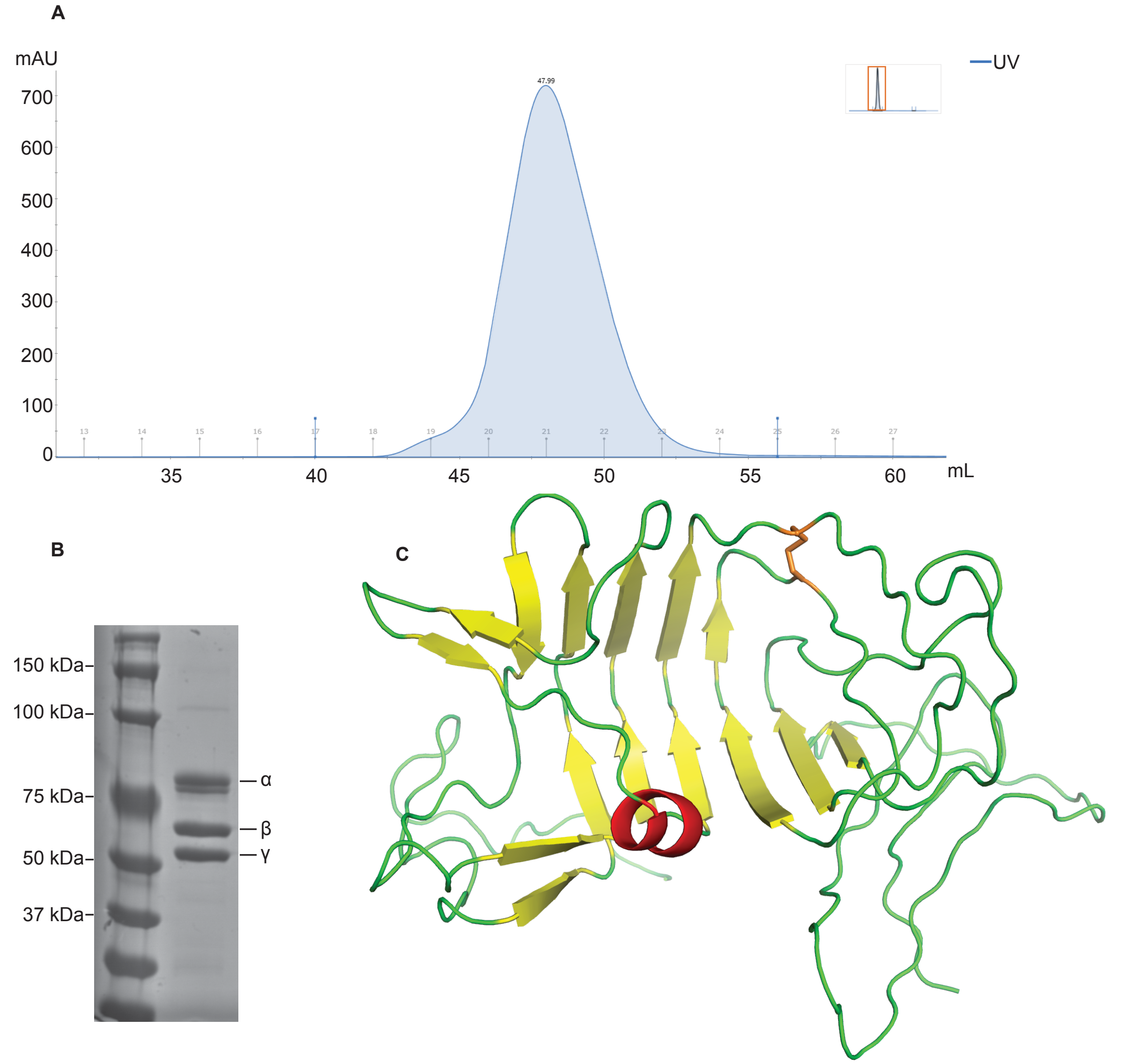


**Supplementary Figure 8. Fibrinogen sample quality, structure model of the αC-domain and superimposition of all fibrinogen structures. (A)** Purification of fibrinogen by S200-Superdex gel filtration with **(B)** reducing SDS 10% PAGE of fractions superimposed **(C)** The predicted structure of the αC-domain. The critical disulphide bond is color in orange.
