## Supplementary figures and images for "Chorography and conformational dynamism of the Soluble Human Fibrinogen in solution"

### SI HDX Table

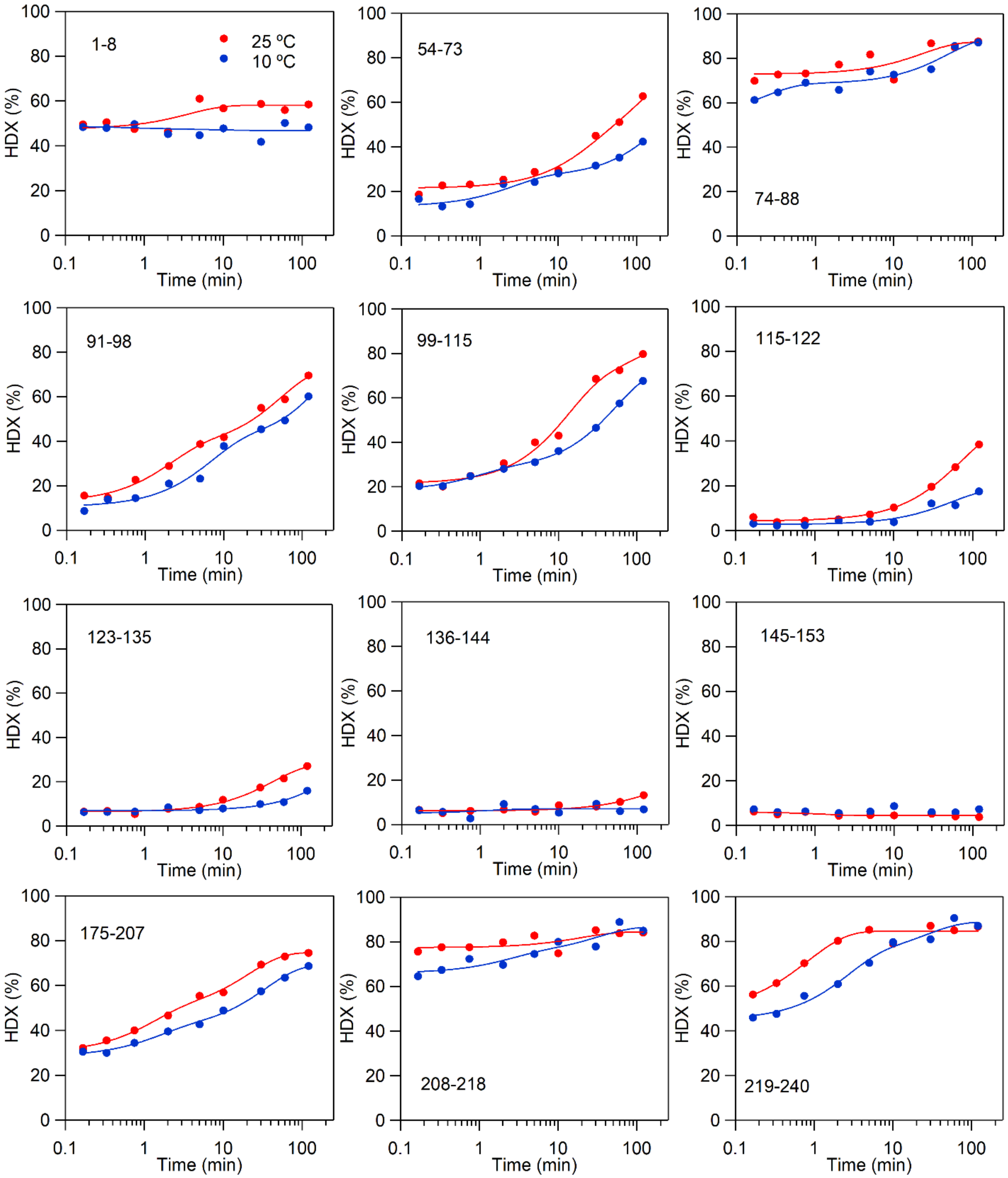

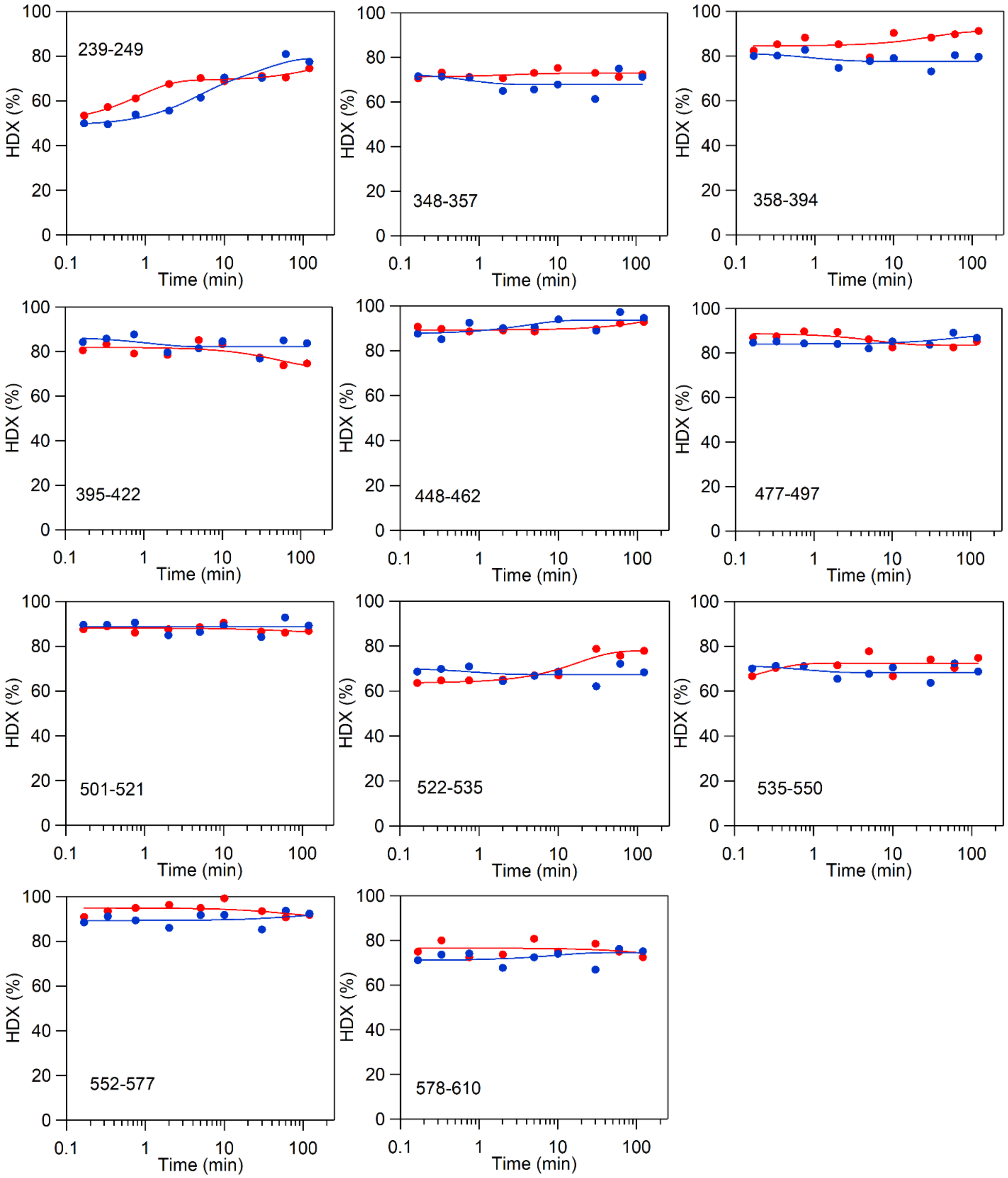

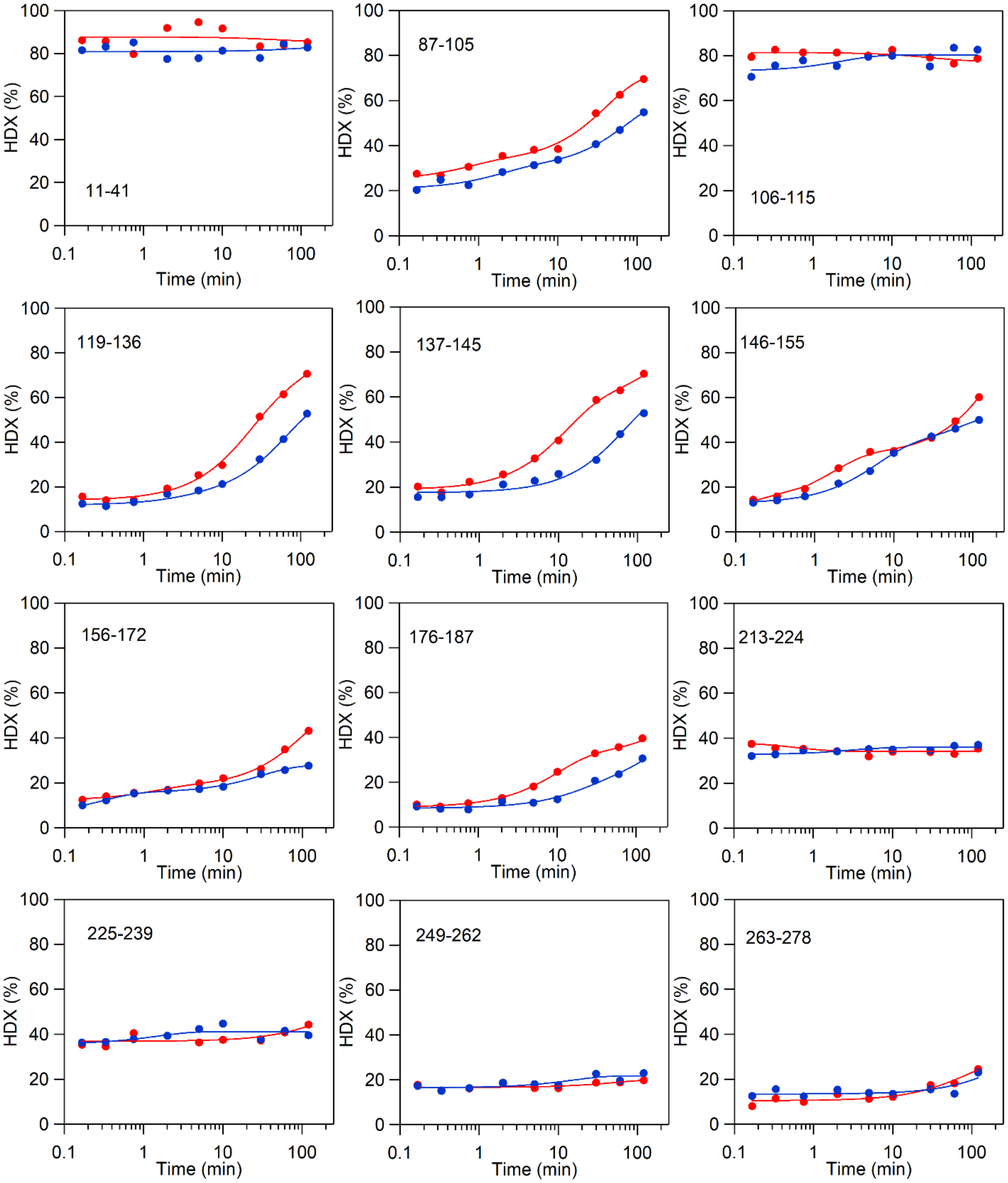

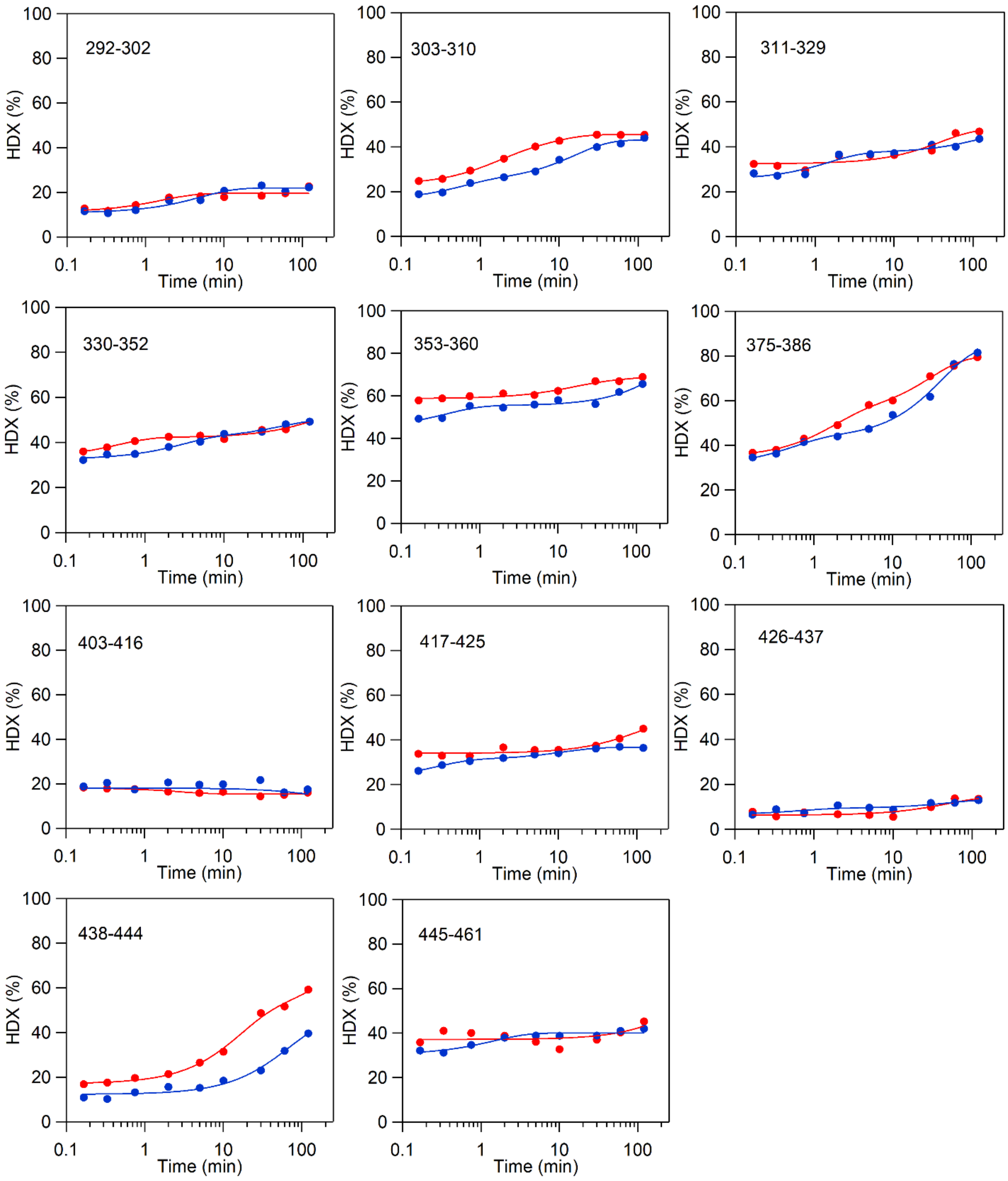

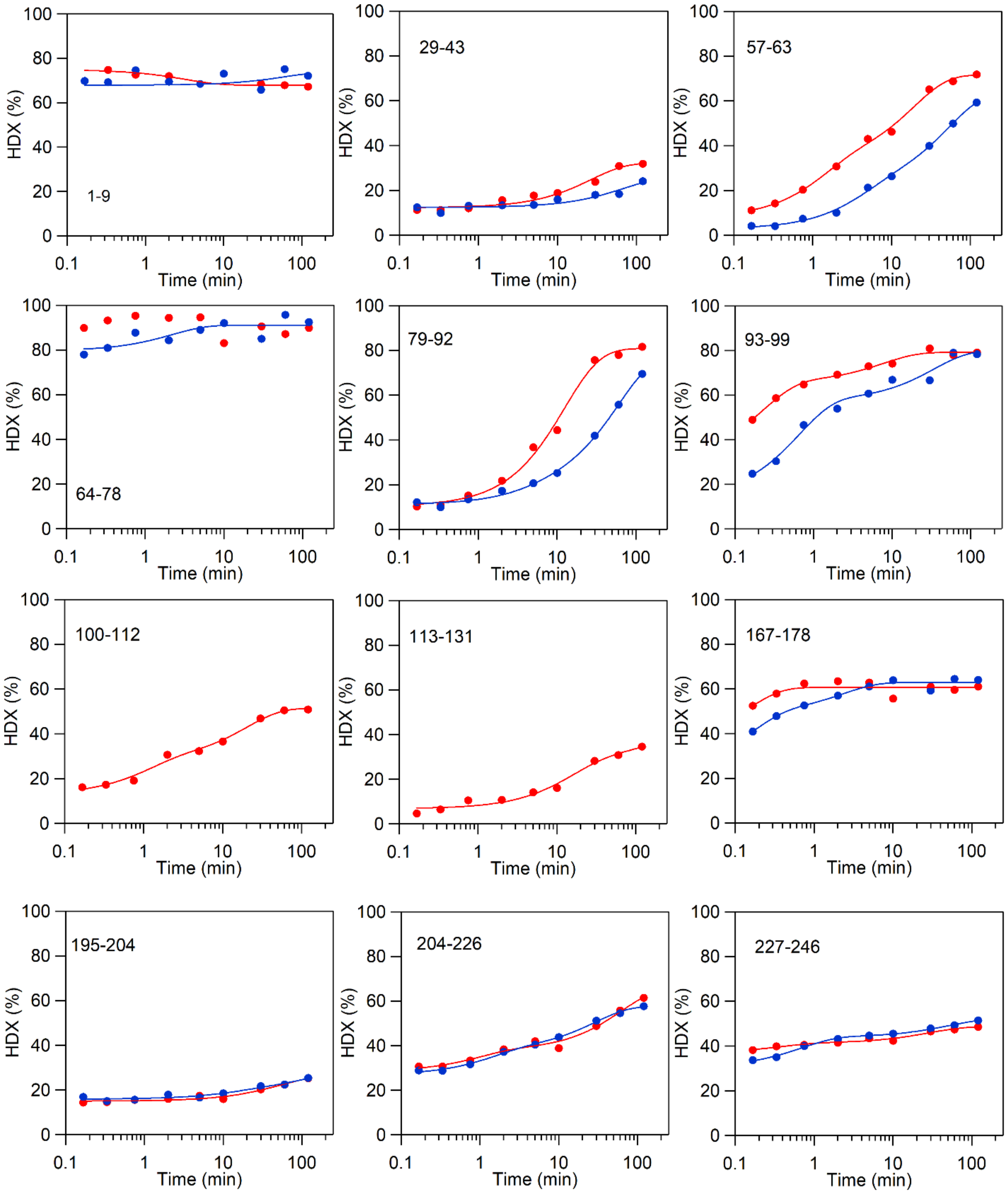

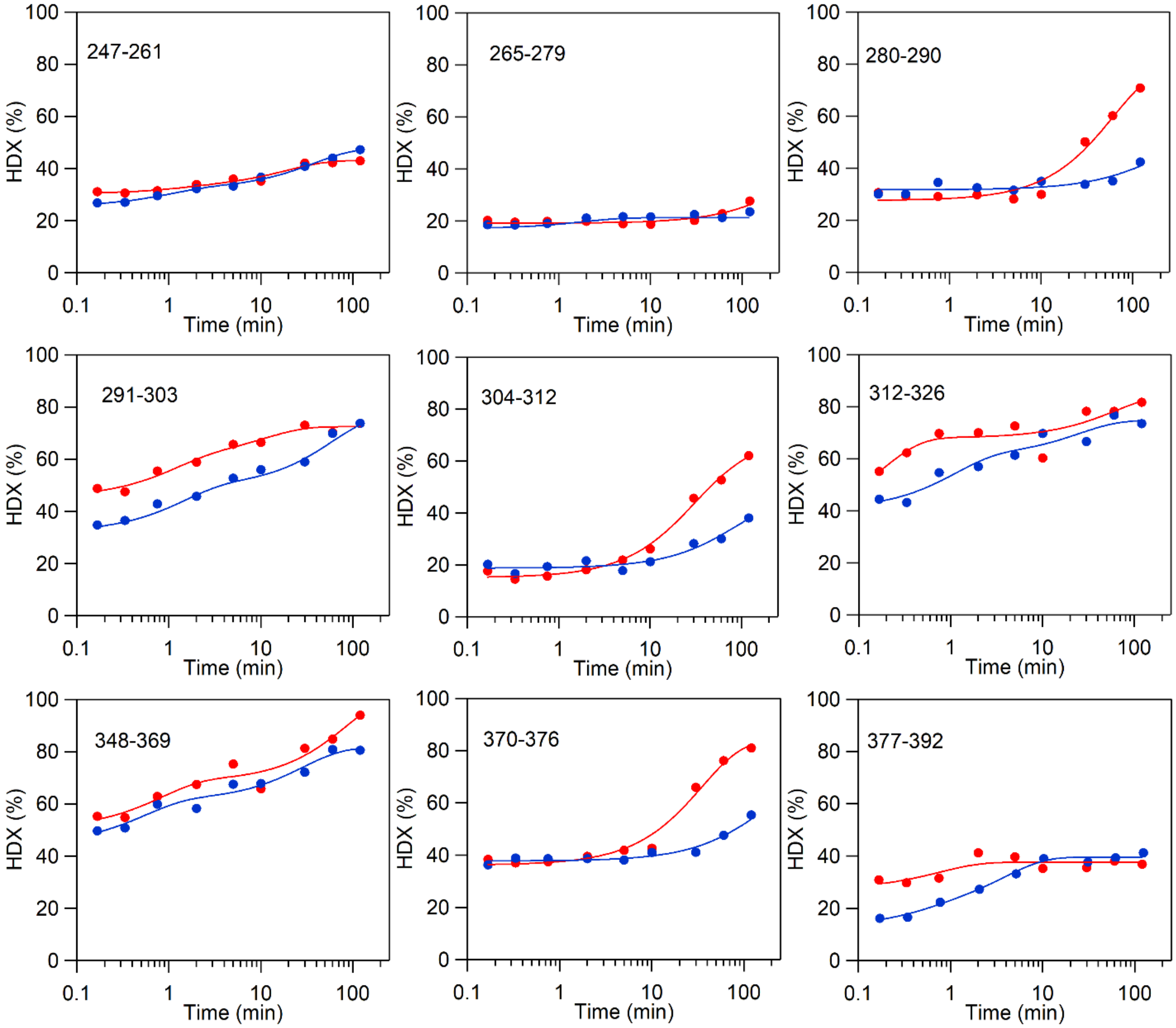
