## Supplementary material for "Chorography and conformational dynamism of the Soluble Human Fibrinogen in solution": SI HDX traces

**Table S1. SAXS sample, data collection, software, and structural parameter information.**

| | **Human Fibrinogen** |
|---|---|
| **(a) Sample details.** | |
| Organism | *Homo sapiens* |
| Source | *Homo sapiens* |
| UniProt sequence ID (residues in construct) | P02671<br>P02675<br>P02679 |
| Extinction coefficient [$A_{280}$, $E^{1\%}$ | 15.1 |
| MW from chemical composition (kDa) | 340 |
| SAXS capillary | |
| Loading concentration ($mg/ml^{-1}$) | 5 |
| Injection volume (μL) | 200 |
| Flow rate (ml/min) | 0.5 |
| Solvent | 20 mM Hepes pH 7.4<br>150mM NaCl |
| Additives | None |
| **(b) SAXS data-collection parameters.** | |
| Instrument/data processing | BNL X9 beamline with 300K PILATUS |
| Wavelength (Å) | 0.918 |
| Camera length (m) | 3.5 |
| q measurement range ($Å^{-1}$) | 0.005-3.19 |
| Absolute scaling method | Comparison with scattering from 1 mm pure $H_2O$ |
| Normalization | To transmitted intensity by beam-stop counter |
| Exposure time | Continuous 2 s data-frame measurements of capillary flow |
| Sample temperature ($^oC$) | 20 |
| **(c) Software employed for SAXS data reduction, analysis and interpretation.** | |
| SAXS data reduction | I(q) versus q and solvent subtraction using BNL in-house software pyXS{Yang, 2020 #1248} |
| Extinction coefficient estimate | ProtParam |
| Basic analyses: Guinier, P(r), *V*P | PRIMUS from ATSAS 2.7.2 {Franke, 2017 #604} |
| Shape/bead modeling | GASBOR {Franke, 2017 #604} via ATSAS online (https:// www.embl-hamburg.de/biosaxs/atsas-online/) |
| Atomic structure modelling | CORAL {Franke, 2017 #604} via ATSAS online (https:// www.embl-hamburg.de/biosaxs/atsas-online/) |
| Three-dimensional graphic model representations | *PyMOL* v.1.8.0.6{Schrodinger, 2015 #856} |
| **(d) Structural parameters.** | **Human Fibrinogen** |
| Guinier analysis | |
| *I*(0) | 53.49 ± 1.651 |
| *R*g (Å) | 135.7 ± 2.688 |
| *P(r)* analysis | |
| *I*(0) | 53.48 |
| *R*g (Å) | 134.5 |
| *d*max (Å) | 448 |
| q range ($Å^{-1}$) | 0.005–0.5 |

**Table S2.** HDX data summary for Fgn

| Data sets | Wild-type human fibrinogen |
|---|---|
| **HDX Reaction Details** | Labeling conditions: 5 μM protein, 90% $D_2O$, 20 mM HEPES pD = 7.4, 150 mM NaCl, 0.5 mM EDTA; corrected pD = $pH_{read}$ +0.4.<br>0s timepoint was collected with 10 μM protein, 20 mM HEPES, 150 mM NaCl, pH = 7.4.<br>Two temperatures: 10 and 25˚C |
| **HDX Time Course** | 10 time points (0, 10, 20, 45, 60, 180, 600, 1800, 3600, 7200 s) for each temperature |
| **HDX Controls** | Maximally-labeled control |
| **Back Exchange: Average, Interquartile** | x%, x% |
| **Number of peptides** | X total; x selected for analysis |
| **Sequence coverage** | x% of peptides were measurable |
| **Average Peptide Length** | x amides (x-x) |
| **Replicates** | 1 (biological) for each temperature<br>Each time point for each temperature was collected once.<br>Due to the scale and purpose of the experiment, it was impractical to collect multiple replicates at all points. To mitigate systematic errors, each temperature set (10 time points) was collected over three days in a non-sequential order. |
| **Repeatability** | There are no replicates, however, the temperature dependence provides a check for outlying datasets. |
| **Significant Differences in HDX** | In this study, we are interested in identifying and classifying the temperature-dependence HDX properties of human fgn. |
